# Structure of dynein-dynactin on microtubules shows tandem recruitment of cargo adaptors

**DOI:** 10.1101/2022.03.17.482250

**Authors:** Sami Chaaban, Andrew P. Carter

## Abstract

Cytoplasmic dynein is a microtubule motor that is activated by its cofactor dynactin and a coiled-coil cargo adaptor. There is currently limited structural information on how the resulting complex interacts with microtubules and how adaptors are recruited. Here, we develop a cryo-EM processing pipeline to solve the high-resolution structure of dynein-dynactin and the adaptor BICDR1 bound to microtubules. This reveals the asymmetric interactions between neighbouring dynein motor domains and how it relates to their motile behaviour. We find unexpectedly that two adaptors occupy the complex. Both adaptors make similar interactions with the dyneins but diverge in their contacts with each other and dynactin. Our structure has implications for the stability and stoichiometry of motor recruitment by cargos.

## Introduction

Eukaryotic cells use their cytoskeleton and motor proteins to organize their cytoplasm. Cytoplasmic dynein-1 (hereafter dynein) is the primary transporter of cargo towards the minus ends of microtubules and therefore contributes to a multitude of cellular processes [Reck-Peterson et al., 2018]. Dynein is a 1.4 MDa multi-subunit dimer that has an intricate autoinhibition mechanism [Zhang et al., 2017]. To be fully active, it must come together with a 1.1 MDa complex, dynactin, and a coiled-coil cargo adaptor [McKenney et al., 2014; Schlager et al., 2014]. Cargo adaptors that share the ability to bind cargo and activate dynein-dynactin motility are referred to as activating adaptors [Reck-Peterson et al., 2018; Olenick and Holzbaur, 2019]. Cryo-electron microscopy (Cryo-EM) of the active complex has shown how the coiled-coils of these adaptors are poised along dynactin’s filament, linking and positioning dynein in a conformation that relieves it from autoinhibition [Urnavicius et al., 2015; Zhang et al., 2017; Urnavicius et al., 2018].

Previous studies showed some adaptors are able to recruit two dyneins per dynactin, increasing the force and speed of the complex, allowing it to outcompete kinesin in a tug-of-war [Urnavicius et al., 2018; Grotjahn et al., 2018; Elshenawy et al., 2019; Elshenawy et al., 2020; Htet et al., 2020]. For the complex to come together, adaptors make multiple interactions with dynein and dynactin [Urnavicius et al., 2015; Gama et al., 2017; Urnavicius et al., 2018; Lee et al., 2020]. These include sites at the adaptor N-termini that bind to the flexible end of the dynein light intermediate chain (LIC) [Reck-Peterson et al., 2018], interactions of the adaptor coiled-coil with dynactin’s pointed end complex [Urnavicius et al., 2015; Gama et al., 2017], and multiple interactions with the dynein heavy chains [Urnavicius et al., 2015; Urnavicius et al., 2018].

Dynein motility is driven by an enzymatic cycle in its motor domains, which connect the dynein-dynactin-adaptor complex to microtubules. Each motor comprises a ring of six AAA+ domains, which, unlike other AAA+ family proteins, are non-equivalent to each other [Roberts et al., 2009]. The nucleotide state of the first AAA+ domain (AAA1) is communicated over a long distance to the microtubule binding domain through a coiled-coil stalk [Gibbons et al., 2005]. The ATPase cycle drives transitions between bent and straight conformations of the linker domain that define dynein’s powerstroke [Schmidt and Carter, 2016]. This communication is gated by the nucleotide states of AAA3 and AAA4 [Bhabha et al., 2014; DeWitt et al., 2015; Nicholas et al., 2015; Liu et al., 2020], and may be affected by interactions between neighbouring dyneins in the assembled complex [Elshenawy et al., 2019].

Although we have structural information on the overall architecture of the dynein-dynactin complex [Urnavicius et al., 2015; Chowdhury et al., 2015; Urnavicius et al., 2018; Grotjahn et al., 2018; Lau et al., 2021], a high-resolution structure of the full complex on microtubules is required to understand how interactions between the motors relate to the way dynein walks and what specifies adaptor recruitment. To achieve this, we took a single particle cryo-EM approach. We used dynein, dynactin and the adaptor BICDR1 (also known as BICDL1, but hereafter referred to as BICDR) which we sparsely-decorated on microtubules in the presence of the ATP-analogue AMPPNP. To overcome the challenge of picking and aligning complexes in the presence of strong tubulin signal, we developed a pipeline to subtract the microtubules from the micrographs. The resulting structures resolved the four motor domains showing that they exist in two major conformations. Averaging the motor domains provided sufficient resolution to reveal nucleotides, and showed that AMPPNP bound in AAA3 locks ADP at the main hydrolysis site (AAA1). Surprisingly, the structure also revealed the presence of a second BICDR adaptor. Taken together, our data suggests that a secondary cargo binding site is a common feature of motile complexes.

## Results

### A microtubule-subtraction pipeline to solve microtubule-bound structures

To solve the structure of the microtubule-bound complex, we purified and reconstituted dynein, dynactin, and BICDR *in vitro*. We chose BICDR due to its robust ability to form dynein-dynactin complexes [Urnavicius et al., 2018]. To keep the complex bound to microtubules, we used the ATP analogue AMPPNP [Grotjahn et al., 2018]. By pelleting the bound dynein-dynactin-BICDR with taxol-stabilized microtubules, we could obtain high contrast cryo-EM images (Figure S1A). The resolution of a previous cryo-ET structure of dynein-dynactin on microtubules was limited by the number of particles due to the low throughput of cryo-ET data collection [Grotjahn et al., 2018]. We therefore used a single-particle approach to obtain a large number of particles (628,033), which enabled high resolution structure determination. The sparse microtubule decoration meant that typical microtubule reconstruction techniques [Zhang and Nogales, 2015; Cook et al., 2020] could not be used to solve the structure. Furthermore, applying conventional single particle analysis techniques was unsuccessful due to the dominating signal of the microtubule, as was observed previously by cryo-ET [Grotjahn et al., 2018] and in cryo-EM of axonemal dyneins bound to microtubule doublets [Ma et al., 2019; Walton et al., 2021; Kubo et al., 2021; Rao et al., 2021]. To overcome this, we developed a pipeline (described below) to subtract the microtubule signal from the micrographs (Figure 1A), which ultimately allowed us to successfully align the complexes (Figure 1B).

**Figure 1.**
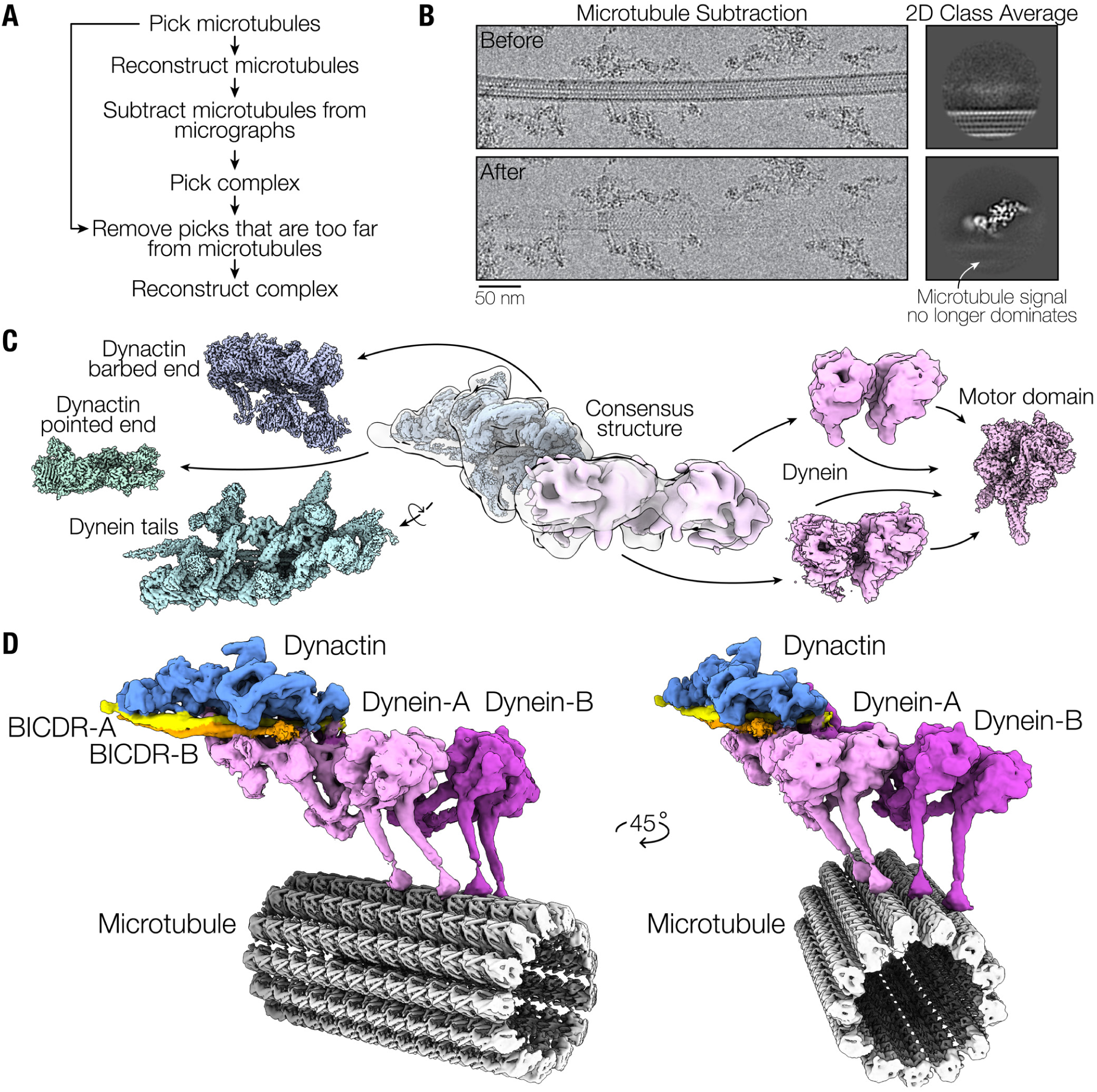
Cryo-EM structure of dynein-dynactin-BICDR on microtubules. (A) Overview of the processing pipeline to subtract microtubules from cryo-EM images in order to thoroughly pick and accurately align dynein-dynactin-BICDR complexes. (B) An example of a microtubule before and after microtubule subtraction. Before subtraction, the microtubule signal dominates, as seen from 2D class averages on the right where the complex is blurred. After subtraction, the complex can be successfully aligned. (C) A gallery of locally processed density maps, starting with the consensus structure in the middle. (D) Composite density map of dynein-dynactin-BICDR overlaid on the reconstructed microtubule.

We first used crYOLO to automatically pick the microtubules from the micrographs [Wagner et al., 2020] and solved the structure of each of the 11 to 15 protofilament number microtubules (Figure S1B) [Cook et al., 2020]. We used the 3D density to computationally subtract the signal from the micrographs in Relion (Figure 1B) [Kimanius et al., 2021]. The subtracted micrographs were then used to pick dynein-dynactin-BICDR complexes in crY-OLO [Wagner et al., 2019]. This allowed us to recover complexes that were otherwise obscured by the overlapping microtubule signal. We then used the microtubule coordinates to remove particles that were deemed too far from a microtubule to be bound (Figure S1C). Taking a single particle approach, we solved the overall structure of the complex and used local refinements to improve the resolution, which provided high resolution coverage of the majority of the complex (Figure 1C, Figure S2).

### Ultrastructure of the dynein-dynactin-BICDR complex on microtubules

Our composite dynein-dynactin-BICDR structure shown in Figure 1D shows the two dynein dimers (dynein-A and B) stacked side-by-side. Each dynein heavy chain (dynein-A1, A2, B1, and B2) contains a motor domain contacting the microtubule and a tail extending towards dynactin (Figure S3A). Each tail binds a LIC and an intermediate chain (IC) (Figure S3B), and binds in grooves in dynactin’s Arp1 filament (Figure S3C). The main BICDR coiled-coil spans dynactin from the barbed end to the pointed end (hereafter BICDR-A) (Figure 1D, Figure S3D). Unexpectedly, a second coiled-coil is present in the structure, which we assign to a second BICDR (hereafter BICDR-B), described below.

In our structure, a previously unobserved density is present that contacts the tail of dynein-A1 (Figure S3E). The density connects to the shoulder domain of dynactin, close to where the ∼75-nm long arm of the dynactin component p150^Glued^ emerges [Urnavicius et al., 2015]. Therefore, the extra density may belong to the globular segment of p150^Glued^ known as the Inter-Coiled Domain (ICD) [Urnavicius et al., 2018]. This suggests that p150^Glued^ may be able to transiently dock onto the tail and motor domain of dynein-A1.

### AMPPNP in AAA3 locks ADP in AAA1

In dynein motor domains, ATP binding and hydrolysis in AAA1 drives the mechanochemical cycle [Schmidt and Carter, 2016]. This cycle can be gated by ATP binding to AAA3, which induces pauses during movement [DeWitt et al., 2015]. A crystal structure of *S. cerevisiae* dynein in the presence of AMPPNP suggests that this inhibition results from the ATP state of AAA3 preventing AAA1 from undergoing hydrolysis [Bhabha et al., 2014]. Given the direct role of microtubules in regulating dynein’s catalytic cycle [Kon et al., 2009], we wanted to re-examine the question of AAA3 gating in the context of our microtubule-bound complex. All of the motor domains in our structure are locked in a post-powerstroke state, where the linker is straight (Figure S4A). We were therefore able to average them to produce a structure with sufficient resolution to reveal side chains and nucleotides (Figure 2A, Figure S2).

**Figure 2.**
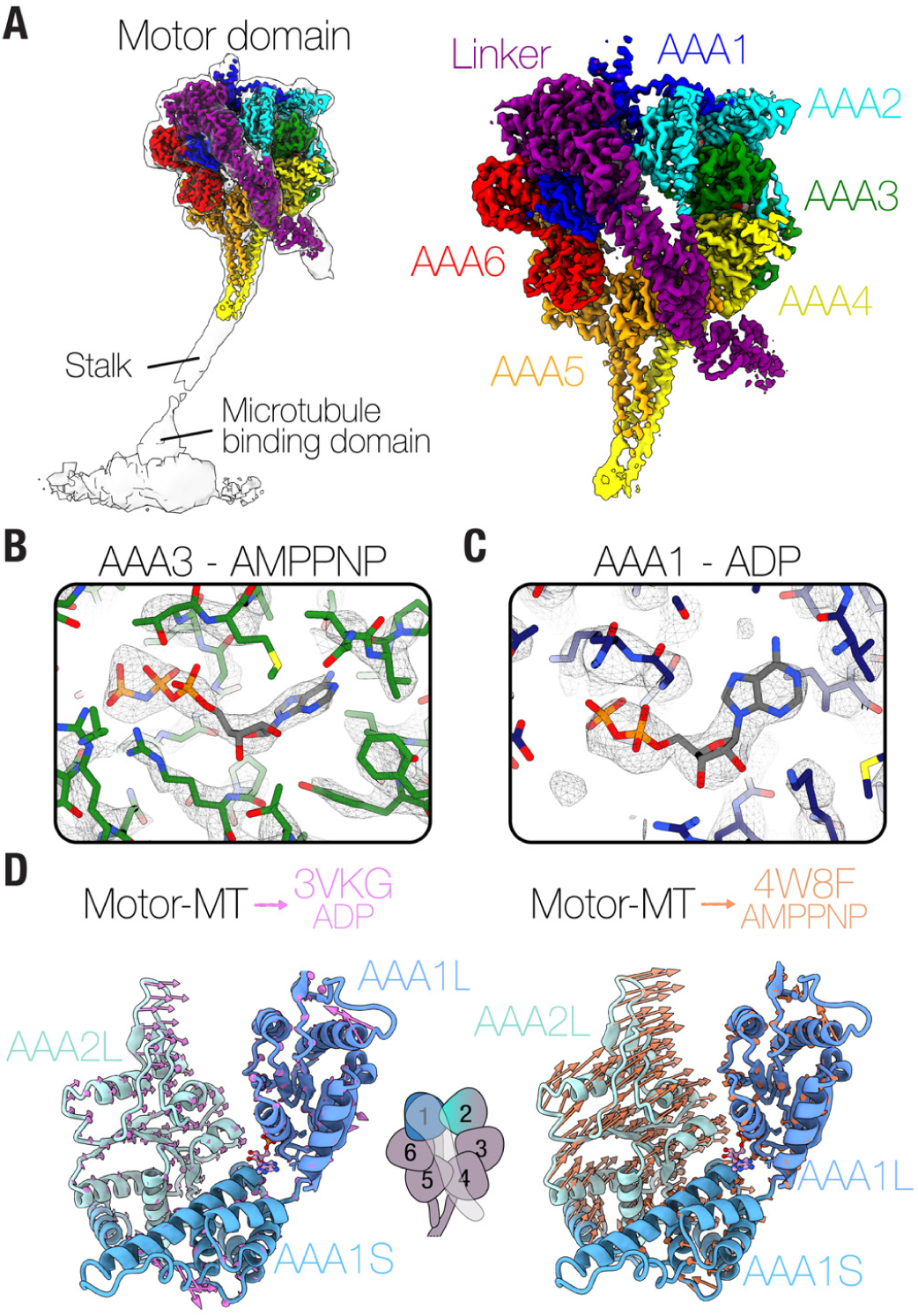
Structure of the dynein motor domain with a trapped ADP molecule in AAA1. (A) The density map of the dynein motor domain placed in the context of a lower resolution refinement of the full motor (left). The subdomains are highlighted on the right. (B) The density in the nucleotide pocket at AAA3 with an AMPPNP molecule docked. (C) The density of the nucleotide pocket at AAA1 with an ADP molecule docked. (D) The domain movements in the nucleotide pocket of AAA1 represented by arrows after alignment of our structure (Motor-MT) to the crystal structure of *D. discoideum* dynein-ADP (left; PDB: 3VKG) [Kon et al., 2012] and *S. cerevisiae* dynein-AMPPNP (right; PDB: 4W8F) [Bhabha et al., 2014].

Overall, the structure is similar to the previous AMPPNP crystal structure [Bhabha et al., 2014]. The linker spans across one face of the AAA+ ring, and is docked onto AAA5 and AAA2 (Figure S4B, S4C). Our structure also contains the C-terminal domain, which is on the opposite face of the ring and is absent in *S. cerevisiae* motor domains (Figure S4D). The density for the nucleotide in AAA3 is consistent with AMPPNP (Figure 2B), and the AAA3 nucleotide pocket is closed (Figure S4E). Surprisingly, however, despite incubating the complex in saturating amounts of AMPPNP, the density at AAA1 is consistent with ADP (Figure 2C). In agreement with this, the conformation of the nucleotide pocket in AAA1 matches the crystal structure of ADP-dynein from *D. discoideum*, where it is more open than AMPPNP-dynein (Figure 2D) [Kon et al., 2012; Bhabha et al., 2014]. Our structure does not have a mutation that blocks hydrolysis in AAA1 that was present in the AMPPNP crystal structure, which may explain the discrepancy in the nucleotide. In summary, our microtubule-bound dynein structure suggests that rather than blocking hydrolysis of AAA1, AMPPNP in AAA3 traps AAA1 in an ADP state.

### Interactions of the dynein heavy chain

Classification of our dynein-dynactin complexes revealed that there are two main conformations of the motor domains: one where the they are aligned next to each other (“aligned”) (Figure 3A), and one where dynein-A is behind dynein-B (“staggered”) (Figure 3B). In the aligned state, the four stalks run parallel to each other, as observed previously by cryo-ET [Grotjahn et al., 2018] (Figure 3A). In the staggered state, however, dynein-A1’s stalk lies behind dynein-A2 (Figure 3B). A key difference between the aligned and staggered states is the interaction between dynein-A and B. In the aligned state, there is no contact between the motor domains of the two dynein dimers, with a distance of ∼25 Å between dynein-A2 and B1 (Figure 3A). In the staggered state, the motor domain of dynein-A2 is docked onto the LIC of dynein-B1 (Figure 3B). This contact is on the linker face of dynein-A2 (Figure 3C, 3D) and involves AAA2 and AAA3 (Figure S5A). On the LIC, the contact involves conserved sites on the Ras-like domain (Figure S5A) [Schroeder et al., 2014].

**Figure 3.**
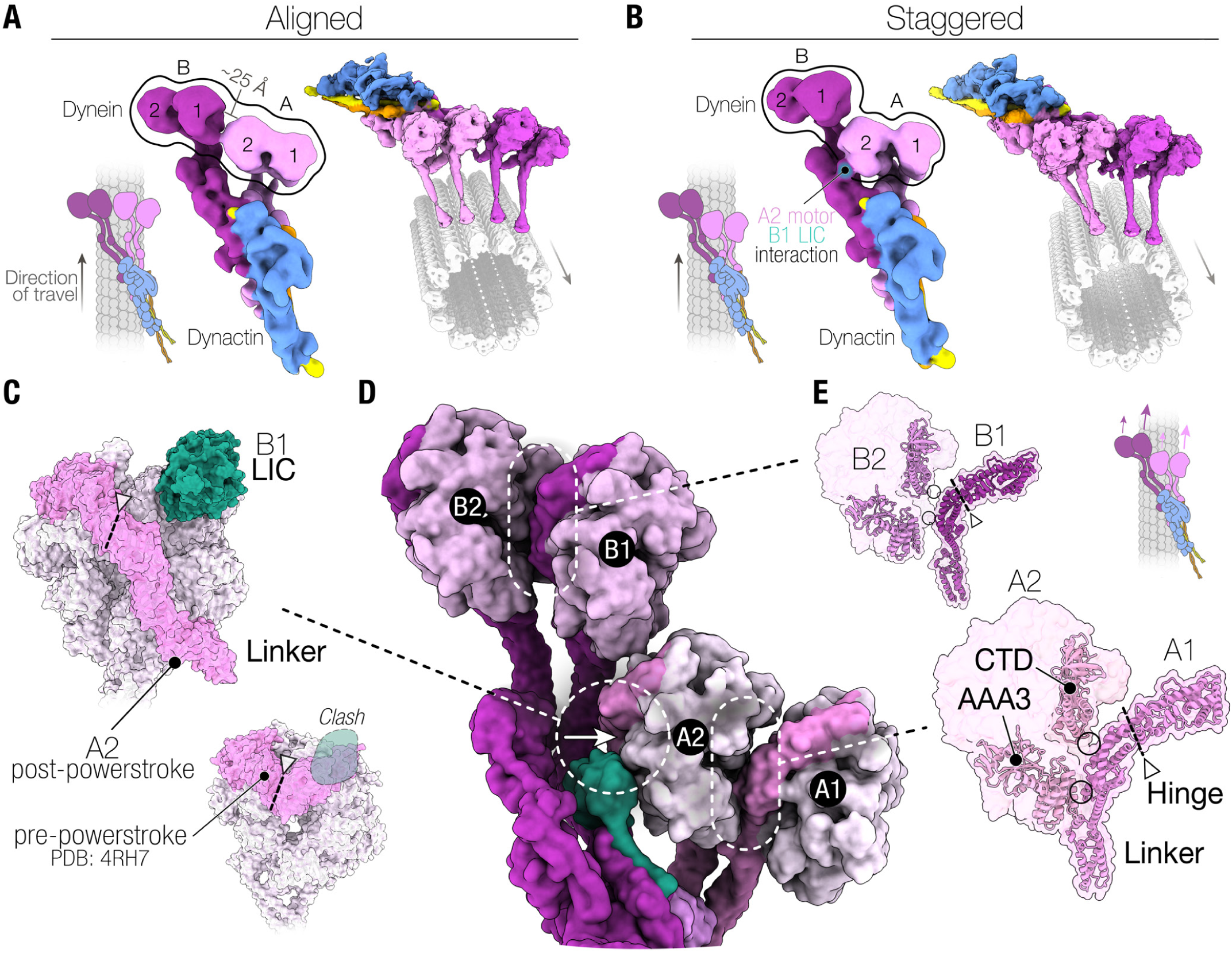
Interactions of the dynein heavy chain. (A) The density map of the aligned state of the dynein motor domains (left). The composite map is overlaid onto the microtubules and viewed from the front (right). (B) The density map of the staggered state of the dynein motor domains (left) and composite map on microtubule (right). (C) Molecular surface representation of the motor domain of dynein-A2 and the LIC of dynein-B1. The bottom right shows the pre-powerstroke state, with the linker bent (PDB: 4RH7) [Schmidt et al., 2015]. The bent linker would clash with the LIC. The triangles point to the hinge in the linker (dotted line). (D) A top view of a molecular surface representation of the dynein heavy chains in the staggered state. (E) A molecular surface representation of the motor domains of the left-side monomers and the linker of the right-side monomers overlaid onto a ribbon representation of the AAA3, C-terminal domain, and linker.

To understand how dyneins influence each other, we analysed their interactions with respect to the movements required for stepping. Looking down on the dynein-dynactin complex in its direction of travel (arrows in Figure 3A and 3B), the motor domains A1 and B1 are on the right of A2 and B2, respectively. The forward step of each motor domain involves a detachment from the microtubule followed by a priming stroke, where the linker bends at its hinge [Carter, 2013]. Since the linker is connected to dynein’s tail, this bending results in a movement of the motor domain. In the staggered state, the consequence of docking onto the LIC of dynein-B1 is that dynein-A2 is prevented from bending its linker. This can be seen by comparing to a pre-powerstroke dynein crystal structure [Schmidt et al., 2015], which shows that the linker would clearly clash with the LIC (Figure 3C). Therefore, dynein-A2 in the staggered state is unable undergo a priming stroke.

The interface between the dynein motor domains within each dimer (i.e. dynein-A1 to A2 and B1 to B2) is similar in both the staggered and aligned states (Figure S5B). The linker from the right-hand motor domain (A1/B1) binds at two points on the left-hand ring (A2/B2), namely AAA3 and the C-terminal domain (Figure 3D, 3E). The interactions on the linkers of dynein-A1 and B1 occur on the tail-side of the hinge, and are therefore not expected to hinder their motor domains from moving during a priming stroke. In contrast, the left-hand monomers, dynein-A2 and B2, bind their neighbours through their rings. This likely impedes their motor domain from moving. Therefore, from our staggered and aligned states, dynein is expected to lead off with the right monomer. This may explain the observed right-handed helical pitch of dynein’s trajectory on microtubules [Elshenawy et al., 2019].

To assess the variability in motor domain interactions, we visualized individual dynein-dynactin-BICDR complexes by cryo-ET. In the tomograms, while some complexes lie in the staggered and aligned states described above (Figure S5C), we observed deviations in the positions of individual motor domains (Figure S5D, S5E). For example, in Figure S5D, we see shifts in the position of dynein-A1, whereas in Figure S5E there is a large separation (∼16 nm) between dynein-B1 and B2. Despite this variability, there are clearly stable interactions between the motor domains which likely affect dynein’s stepping behaviour.

### Two adaptors scaffold the complex

Dyneins are linked by their tails to dynactin via a cargo adaptor. Previously it was reported that one adaptor coiled-coil binds along dynactin, with the N-terminus at the barbed end and the C-terminus at the pointed end [Urnavicius et al., 2018]. The overall conformation is retained in our structure of BICDR-A (Figure 4A), which runs the length of dynactin close to dynein-A and B. Surprisingly, the second adaptor, BICDR-B, only covers half the length of the complex, interacting mainly with dynein-A1. The two adaptors meet close to where BICDR-A binds dynein-A1 (Figure 4B), and there is an extensive interaction between the coiled-coils over ∼30 residues before they diverge again to contact the pointed end of dynactin. Mass photometry of BICDR shows a major peak relating to a single coiled-coil (Figure S6A), suggesting that the interaction between BICDRs is stabilized in the complex. The registry of the BICDR coiled-coils can be determined based on a unique tryptophan (Trp166), which presents as a bulky density between the two alpha helices of the coiled-coil (Figure S6B). This shows that there is an offset of ∼47 residues between the interaction sites of BICDR-A and B (Figure 4B, Figure S6C).

**Figure 4.**
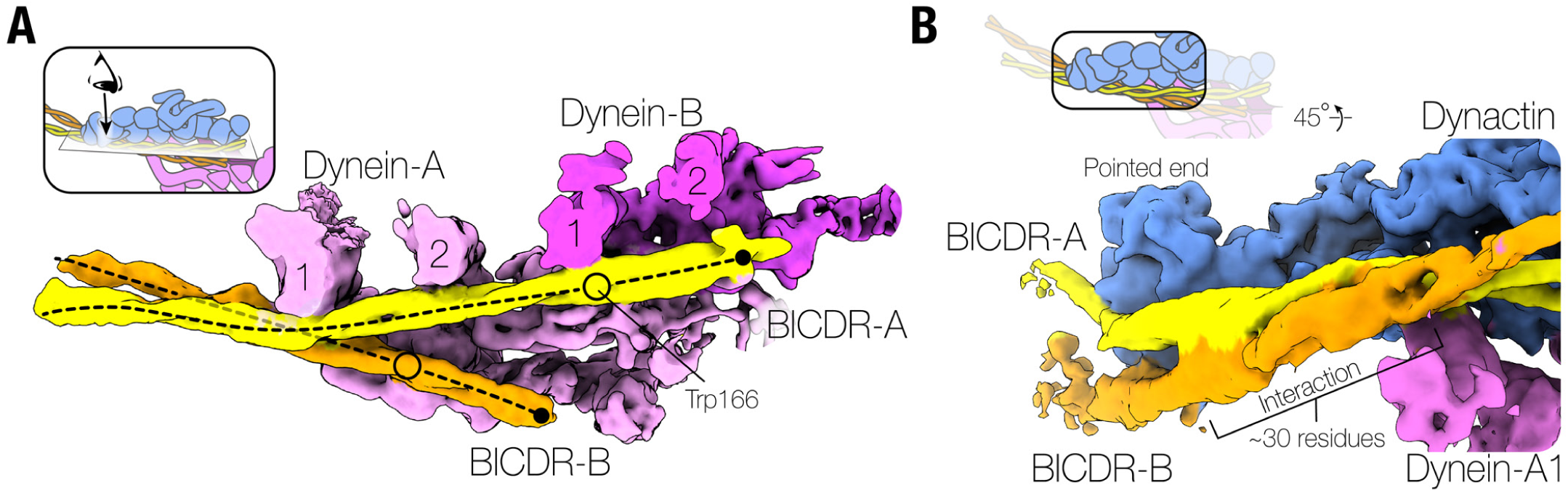
Two BICDRs scaffold the dynein-dynactin complex. (A) A top view of the locally refined density map of the dynactin-dynein-tail region. The density for dynactin is hidden to more clearly see the position of BICDR-B. The position of Trp166 is shown as a black circle for each adaptor. Note that the position of BICDR-B is offset relative to BICDR-A. (B) A side view of the pointed end of dynactin showing where the two BICDRs overlap near dynein-A1 before diverging.

To address whether two adaptors are present in the previous structures of dynein-tail/dynactin/BICDR [Urnavicius et al., 2018], we revisited the data using new 3D classification tools in cryoDRGN [Zhong et al., 2021]. We found density for the second adaptor in a subset of particles (∼5%) (Figure S6D). This suggests that in our new structure, which has almost 100% occupancy of both adaptors, the presence of microtubules stabilizes the binding of BICDR-B.

Two coiled-coils were seen previously at the pointed end of dynactin in a structure with the adaptor Hook3 [Urnavicius et al., 2018; Lau et al., 2021]. However, it was not possible to determine if they belonged to two separate adaptors or resulted from a folding back of Hook3. We can now clearly assign these coiled-coils to two Hook3s (Hook3-A and Hook3-B) using our new classifications (Figure S6E). Taken together, our analysis suggests that the presence of two adaptors may be found in other dynein-dynactin complexes.

### Adaptor motifs define how two BICDRs bind asymmetrically to dynein-dynactin

BICDR interacts with dynein and dynactin via conserved adaptor motifs. These include (1) a CC1 box (AAXXG), which interacts with the LIC, (2) a CC2 box, which interacts with the dynein heavy chain, and (3) a Spindly motif, which interacts with dynactin’s pointed end [Gama et al., 2017; Sacristan et al., 2018; Lee et al., 2018; Celestino et al., 2019].

The CC1 box interacts with a C-terminal alpha helix of the LIC, which extends from the Ras-like domain by a long flexible loop (Figure S3B). Because BICDR is a dimer, the CC1 box region has a LIC binding site on both sides of its coiled-coil [Lee et al., 2018]. Therefore, we expected that the four LIC helices originating from our two dyneins interact with the four LIC binding sites (i.e. two on each adaptor CC1 box). In our structure, we see evidence of LIC binding on both sides of BICDR-A (Figure 5A, Figure S7A). The LIC on the outside face is much less resolved, suggesting it has lower occupancy. 3D classification showed that in some classes, there is density connecting the LIC of dynein-A2 to the CC1 box of BICDR-A (Figure S7B), as previously observed [Urnavicius et al., 2018].

**Figure 5.**
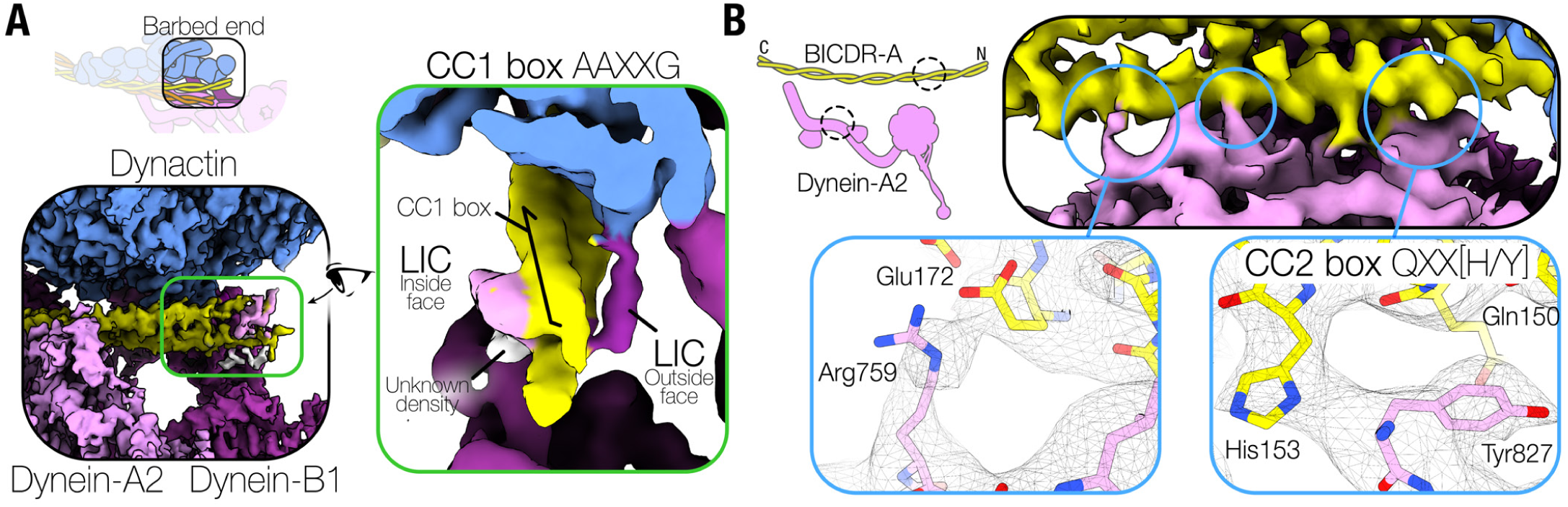
CC1 box and CC2 box interactions of BICDR-A. (A) A close up of the barbed end of dynactin where the N-terminus of BICDR-A is visible (left). A front view of the coiled-coil, filtered to a lower resolution to show densities on both sides of the CC1 box (right). (B) A close up of the interaction between BICDR-A and dynein-A2, highlighting the CC2 box conserved residues (Gln150/His153) and the C-terminal Glu172.

The CC1 box of the second adaptor, BICDR-B, is less defined due to its increased flexibility. However, in some 3D classes, we also observed connecting density from the LIC of dynein-A2 (Figure S7C). This suggests that the LIC helix from dynein-A2 can contact either adaptor. We could not find connecting density to the outside face of the CC1 boxes, probably due to their flexibility or partial occupancy. However, due to their proximity, they are likely accommodated by LICs from dynein-B (contacting BICDR-A) and dynein-A1 (contacting BICDR-B). In summary, the dynein-A2 LIC is a major contributor to BICDR-A and BICDR-B CC1 box interactions. This disfavours our original hypothesis that each adaptor binds the LICs of a separate dynein.

Dynein-A2 docks onto BICDR-A at a site C-terminal to the CC1 box. Previously, another motif implicated in complex formation, the CC2 box (QXX[H/Y]) [Sacristan et al., 2018], was predicted to reside at this interface [Lau et al., 2021]. Our structure now shows that the highly conserved Gln150 and His153 of this motif interact with Tyr827 of dynein-A2 (Figure 5B). Additionally, we can resolve interactions between dynein-A2 and BICDR-A at residues Glu165 and Glu172 (Figure 5B). Based on our predicted registry of BICDR-B, its CC2 box contacts dynein-A1 in a similar way (Figure S6B). Therefore, although the coiled-coils of the two adaptors have distinct positions in the complex, they are both anchored at their N-termini by CC2 box interactions with the monomers of dynein-A.

The C-termini of BICDR-A and BICDR-B diverge in the way they interact with the pointed end of dynactin (Figure 6A). Previous work analysing the binding of different activating adaptors to the pointed end identified four distinct sites of interaction [Lau et al., 2021]. BICDR was reported to bind site 1 (p62 disordered loop), site 2 (p62 saddle region), and site 4 (p25 β-helical fold). In our structure, BICDR-A interacts similarly at site 4 (Figure 6A), but the interactions at site 1 and site 2 are shifted to lower positions on p62, resulting from a downward rotation of the coiled-coil at site 1 (Figure S8A). Unlike BICDR-A, BICDR-B interacts with the pointed end at site 3 (on a loop extending from p25) (Figure 6A).

**Figure 6.**
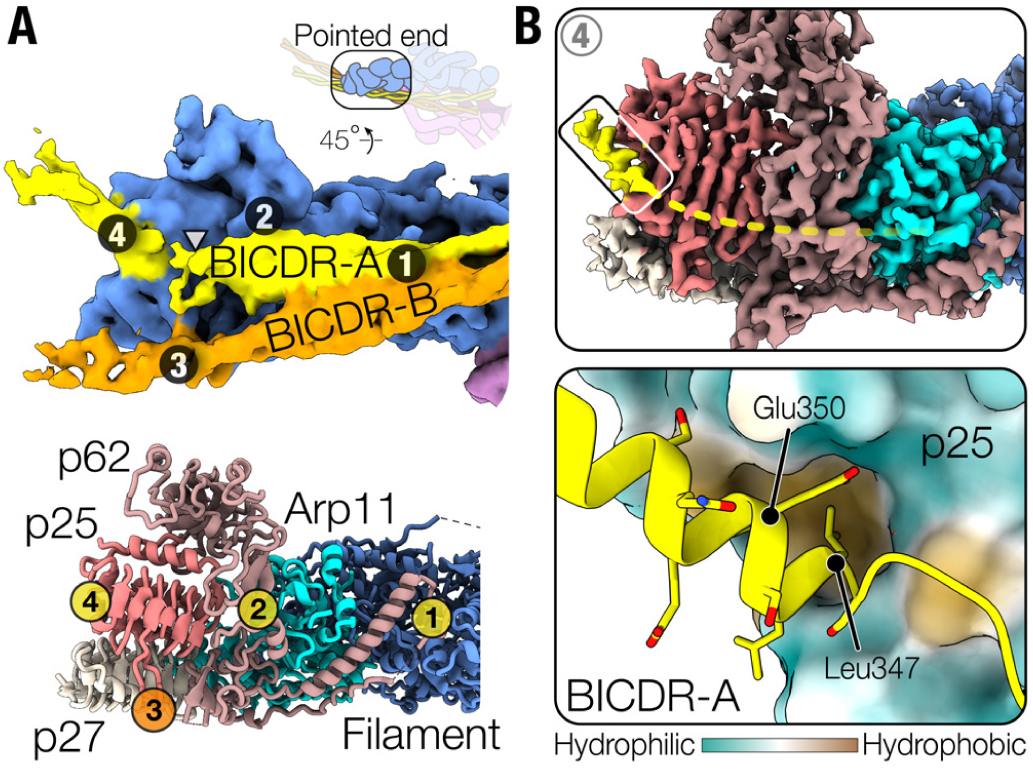
Pointed end interactions with two adaptors. (A) The density map of the pointed end showing how BICDR-A (yellow) and BICDR-B (orange) interact at distinct sites (top), which are numbered 1 to 4 (bottom) [Lau et al., 2021]. The triangle points to the break in the coiled-coil density. (B) The density map of the pointed end showing the ordered alpha helical density at site 4 (top) where the Spindly motif of BICDR-A resides. The N-terminal density is less resolved (dotted yellow line). A molecular surface representation of the pointed end coloured by hydrophobicity (orange is hydrophobic, teal is hydrophilic) (bottom) shows the Spindly motif binding in a hydrophobic pocket in p25.

The pointed end is the site where the Spindly motif (LΦXEΦ, where Φ denotes an aliphatic or aromatic side chain), which is found in many activating adaptors, interacts [Gama et al., 2017]. Our model places it at site 4, as previously predicted [Lau et al., 2021]. Although the alpha-helical density docked onto site 4 is the most ordered part of the BICDR-A C-terminus (Figure 6B), the structure suggests that there is a break in the coiled-coil preceding it (Figure 6A). Therefore, to provide support for our model, we ran an Alphafold prediction [Evans et al., 2021] of the pointed end BICDR complex (Figure S8B). The resulting prediction shows high confidence for the Spindly motif interacting with site 4 as well as a preceding break in the coiled-coil, consistent with the density. Interestingly, other cargo adaptors may also have coiled-coil breaks preceding their Spindly motifs (Figure S8C), as recently predicted [d’Amico et al., 2022]. Our structure now shows that the highly conserved Spindly motif residue Leu347 sits in a hydrophobic pocket at the edge of p25 (Figure 6B). The interaction is further stabilized by the adjacent Leu348 and the highly conserved Glu350 interacting with Ser31 and Asn20 of p25, respectively. Due to the offset between BICDR-A and BICDR-B, the Spindly motif of BICDR-B does not contribute to its pointed end interaction.

In summary, whereas the N-terminal motifs (CC1 and CC2 boxes) of the two adaptors bind similarly with dyneins, the C-terminal interactions with the dynactin pointed end are distinct, leading to an asymmetry in the two adaptors (Figure S8D).

## Discussion

### Solving structures of flexible complexes on filaments

Solving the structure of proteins decorating microtubules poses unique challenges, especially when the protein is sparsely and randomly decorated. The microtubule signal dominates the alignments and makes it difficult to thoroughly pick all the proteins. Previously, the problem of the dominating microtubule signal was overcome by subtracting the microtubule from picked particles [Ma et al., 2019; Walton et al., 2021; Kubo et al., 2021], but this only works well when the proteins decorate the microtubule as a regular array. Here, we took the approach of reconstructing the microtubules and subtracting them from the raw micrographs, revealing obscured particles and eliminating the influence of the microtubule signal. An alternative approach was recently reported, where 2D averages were calculated along individual microtubules and used to subtract them from micrographs [Chai et al., 2022], although so far this has only been used on regularly decorated samples. These two approaches open the door to solving structures of other sparsely decorated complexes on cytoskeletal filaments.

### Dynein’s high affinity state is regulated by AAA3

The dynein family of motors includes axonemal dyneins, cytoplasmic dynein-2, and cytoplasmic dynein-1, discussed here. In some family members, including cytoplasmic dynein-2, AAA3 only binds ADP [Schmidt et al., 2015]. In other dyneins, AAA3 is competent for ATP hydrolysis [Gleave et al., 2014]. In the case of cytoplasmic dynein-1, ATP binding at this site results in pausing on the microtubule [DeWitt et al., 2015]. The predominant model of how ATP at AAA3 blocks the catalytic cycle is based on the crystal structure of the AMPPNP dynein motor domain, where AMPPNP was found in both AAA1 and AAA3 [Bhabha et al., 2014]. This, along with molecular dynamics simulations [Goldtzvik et al., 2018], suggested that pauses occur due to the inability of AAA1 to hydrolyse ATP. In contrast, our full-length dynein structure, which is on microtubules and does not contain any mutations in AAA1, suggests that when AAA3 binds AMPPNP, AAA1 binds ADP. We therefore propose a model where ATP binding in AAA3 locks the motor domain in a high microtubule affinity, post-powerstroke state by preventing ADP release from AAA1. In addition to causing pauses, this likely affects the binding of Lis1, a dynein regulator that depends on the nucleotide state of AAA3 [DeSantis et al., 2017].

### Asymmetric restraints on the dynein motors

Our analysis of the interaction between the four motor domains within the two-dynein complex suggests that there’s a remarkable asymmetry between them. Dynein-A1 and B1 appear free to undergo a priming stroke whereas dynein-A2 and B2 are constrained. Although our complex is in an AMPPNP-inhibited state, it is striking that to step forward, the right-hand motors (dynein-A1 and B1) appear likely to move first. This asymmetry may explain the primarily right-handed pitch of dynein-dynactin complex trajectories on microtubules [Elshenawy et al., 2019]. More rarely, left-handed trajectories were observed. This is perhaps explained by the heterogeneity of the position of the motor domains, as seen by our cryo-ET analysis of individual complexes. In the presence of ATP, this flexibility is likely to be higher. In summary, the asymmetry we observe may contribute to the coordinated movement of the motor domains.

### Motifs define how adaptors bind in pairs

Cargo adaptors are classified based on the identity of their LIC-binding site, which includes the previously mentioned CC1 box family (e.g. BICDR, Spindly, BICD2, and TRAK1), as well as the Hook domain family (e.g. Hook1 and Hook3), the EF-hand family (e.g. RAB11FIP3 and Cracr2a), and the RH1 domain family (e.g. RILP and JIP3) (Figure S8C) [Reck-Peterson et al., 2018; Olenick and Holzbaur, 2019]. In light of our structure, an important question is which of these adaptors bind to dynein-dynactin in pairs. In BICDR, both adaptors interact with dynein via their CC1 and CC2 boxes. They bind to each other in an offset manner and interact with different sites at dynactin’s pointed end. A second example is the adaptor Hook3, where our new 3D classifications of previous data [Urnavicius et al., 2018] clearly show the presence of two adaptors. The Hook domain is not resolved but biochemical evidence suggests its interaction with the LIC is important for complex formation [Lee et al., 2018]. Both copies of Hook3 interact with the dynein heavy chain in similar positions to BICDR (Figure S6D, S6E). Interestingly, the density suggests that the interaction between the two Hook3s is different from the two BICDRs. Furthermore, Hook3 and BICDR make different interactions with the pointed end [Lau et al., 2021]. Specifically, whereas BICDR-A contacts sites 1/2/4, Hook3-A binds sites 1/3/4 (Figure S8E). Conversely, whereas BICDR-B contacts site 3, Hook3-B binds site 2. This suggests that there is considerable diversity in how pairs of adaptors interact with the dynein-dynactin complex.

Given the anchoring interaction of the CC2 box in both BICDRs, we wondered if the motif was also present in Hook3, since it was previously only described for the CC1 box family [Sacristan et al., 2018]. A sequence alignment confirms that Hook3 contains the motif, as well as the C-terminal glutamates that we show accompany the interaction (Figure S9A). An Alphafold prediction of the dynein heavy chain and CC2 box region of Hook3 further supports the interaction (Figure S9B).

The BICDR and Hook3 structures suggest that a LIC-binding site and CC2 box are important for recruiting a second adaptor. These motifs have been predicted in several cargo adaptors (e.g. Spindly, TRAK1/2, and HAP1) [Sacristan et al., 2018] (Figure S8C), suggesting that they may also be able to bind in pairs. A further implication is that it may be possible to recruit a mixture of adaptors to the same dynein-dynactin complex. This is especially relevant for those with shorter coiled-coils that lack Spindly motifs (e.g. RILP and JIP3) (Figure S8C), which may hitchhike with longer coiled-coil adaptors.

### Consequences of having a second adaptor

BICDR and Hook3 transport predominantly membrane vesicles [Schlager et al., 2010; Bielska et al., 2014]. It is conceivable that two vesicles could be recruited simultaneously to the complex, one by each adaptor, but their large size suggests that this is unlikely, especially in axons [Foster et al., 2021]. Instead, a single dynein-dynactin complex is probably linked to the membrane by two connections. This bivalent tethering of motors to vesicles may increase the efficiency of motor capture and decrease the probability of release. For small molecules, such as RNAs transported by the related adaptor BicD [McClintock et al., 2018], there is the possibility that dynein-dynactin can carry two cargos at once. This would allow greater capacity for cargo transport.

Another consequence of the second adaptor binding site is that it reinforces the connection between dynein-A and dynactin. This will reduce the dissociation of the dynein-dynactin complex, which is likely to promote long distance movement on microtubules. Since both adaptors primarily contact dynein-A, this stabilization is likely to also occur in adaptors which can recruit a single dynein (e.g. BICD2) [Urnavicius et al., 2018].

Many dynein cargo adaptors can also bind kinesins [Hancock, 2014; Fu and Holzbaur, 2014], which leads to complexes containing opposing motors [Fenton et al., 2021; Canty et al., 2021]. The relative number of each motor may be key to determining the behaviour of cargo movement [Rai et al., 2016; Belyy et al., 2016]. Our observation that two adaptors can bind per dynein-dynactin complex suggests that they may also recruit additional kinesins. We therefore speculate that different adaptors can tune the stoichiometry of the motors that they recruit, providing an added layer of regulation to cargo transport.

## Methods

### Protein preparation

Dynactin was purified natively from pig brains [Urnavicius et al., 2015]. Fresh brains were acquired from a butcher and transported in ice cold PBS, cleaned in homogenization buffer (35 mM PIPES pH 7.2, 5 mM MgSO_4_, 100 μM EGTA, 50 μM EDTA), and flash frozen in liquid nitrogen. Frozen brain pieces were blended and resuspended in homogenisation buffer supplemented with 1.6 mM PMSF, 1 mM DTT, and 4 complete-EDTA protease-inhibitor tablets per 500 mL (Roche). After thawing, the lysate was centrifuged in a JLA 16.250 (Beckman Coulter) at 16,000 rpm for 15 min at 4°C. The supernatant was further clarified in a Type 45 Ti (Beckman Coulter) at 45,000 rpm for 50 min at 4°C. After filtering the supernatant in a Glass Fibre filter (Sartorius) and a 0.45 μm filter (Elkay Labs), it was loaded on a column packed with 250 mL of SP-Sepharose (Cytiva) pre-equilibrated with SP buffer (35 mM PIPES pH 7.2, 5 mM MgSO_4_, 1 mM EGTA, 0.5 mM EDTA, 1 mM DTT, 0.1 mM ATP) using an Akta Pure (Cytiva). The column was washed with SP buffer with 3 mM KCl before being eluted in a linear gradient up to 250 mM KCl over 3 column volumes. The peak around ∼15 mS/cm was collected and filtered in a 0.22 μm filter (Elkay Labs) before being loaded on a MonoQ 16/10 column (Cytiva) pre-equilibrated with MonoQ buffer (35 mM PIPES pH 7.2, 5 mM MgSO_4_, 100 μM EGTA, 50 μM EDTA, 1 mM DTT). The column was washed with MonoQ buffer before being eluted in a linear gradient up to 150 mM KCl over 1 column volume, followed by another linear gradient up to 350 mM KCl over 10 column volumes. The peak around ∼39 mS/cm was pooled and concentrated to ∼3 mg/mL before being loaded on a TSKgel G4000_SWXL_ (Tosoh Bioscience) pre-equilibrated with GF150 buffer (25 mM HEPES pH 7.2, 150 mM KCl, 1 mM MgCl_2_) supplemented with 5 mM DTT and 0.1 mM ATP. The peak at ∼114 mL was pooled and concentrated to ∼3 mg/mL. 3 μL aliquots were flash frozen in liquid nitrogen and stored at −80°C.

Human cytoplasmic dynein-1 with an N-terminal ZZ-TEV tag on the heavy chain was expressed using the baculovirus/Sf9 system [Schlager et al., 2014]. Specifically, we used a construct with mutations in the linker (R1567E and K1610E) to help overcome the autoinhibited conformation [Zhang et al., 2017]. Fresh bacmid DNA was transfected into Sf9 cells at 0.5×10^6^ cells/mL in 6-well cell culture plates using FuGene HD (Promega) according to the manufacturer’s protocol (final concentration 10 μg/mL). After three days, 1 mL was added to 50 mL of 1×10^6^ cells/mL and infected for five days in a shaking incubator at 27°C. The P2 virus was isolated by collecting the supernatant after centrifugation at 4,000 rcf for 15 min and stored at 4°C. For expression, 10 mL of P2 virus was used to infect 1 L of Sf9 cells at 1.5-2×10^6^ cells/mL for 96 hours in a shaking incubator at 27°C. Cells were harvested by centrifugation at 4,000 rcf for 10 min at 4°C, resuspended in cold PBS, and centrifuged again in 50 mL Falcon tubes (Sarstedt). The pellet was flash frozen and stored at −80°C. A pellet from 1 L expression was thawed on ice in 50 mL lysis buffer (50 mM HEPES pH 7.4, 100 mM NaCl, 10% (v/v) glycerol, 0.1 mM ATP) supplemented with 2 mM PMSF, 1 mM DTT, and 1 complete-EDTA protease-inhibitor tablets per 50 mL. Cells were lysed using a 40 mL dounce tissue grinder (Wheaton) with ∼16 strokes. The lysate was clarified at 503,000 rcf for 45 min at 4°C using a Type 70 Ti Rotor (Beckman Coulter). The supernatant was incubated with 3 mL IgG Sepharose 6 Fast Flow beads (Cytiva) pre-equilibrated with lysis buffer for 4 hours in a 50 mL Falcon at 4°C on a roller at a low speed setting. The beads were then applied to a gravity flow column and washed with 150 mL of lysis buffer and 150 mL of TEV buffer (50 mM Tris-HCl pH 7.4, 150 mM KAc, 2 mM MgAc, 1 mM EGTA, 10% (v/v) glycerol, 0.1 mM ATP, 1 mM DTT). The beads were then transferred to a 5 mL centrifuge tube (Eppendorf) and filled up completely with TEV buffer. TEV protease, which was purified in-house using pRK793 [Kapust et al., 2001], was added to a final concentration of ∼60 ug/mL and incubated overnight at 4°C on a roller at a low speed setting. The beads were transferred to a gravity flow column and the flow through containing the cleaved protein was collected in a 15 mL Falcon tube. 8 mL of TEV buffer was added to the beads and collected to maximize recovery. The protein was concentrated to ∼2 mg/mL and loaded onto a TSKgel G4000_SWXL_ column preequilibrated with GF150 buffer supplemented with 5 mM DTT and 0.1 mM ATP. Peak fractions were pooled and concentrated to ∼2.5 mg/mL. Glycerol was added to a final concentration of 10% from an 80% stock made in GF150 buffer. 3 μL aliquots were flash frozen and stored at −80°C.

Full-length mouse BICDR1 with an N-terminal ZZ-TEV tag was purified using baculovirus expression in the same way as dynein described above with the omission of ATP in the GF150 buffer.

Lyophilised porcine tubulin (Cytoskeleton) was resuspended in microtubule buffer (25 mM MES, 70 mM NaCl, 1 mM MgCl_2_, 1 mM EGTA, 1 mM DTT, pH 6.5) to a final concentration of 10 mg/mL (∼90 μM), aliquoted, flash frozen, and stored at −80°C. The use of a MES-based buffer increased the proportion of 13 protofilament microtubules (see below) [Pierson et al., 1978].

### Sample preparation

To polymerize microtubules, tubulin was diluted in microtubule buffer with GTP (MilliporeSigma) such that the final concentration of tubulin was 5 mg/mL (45 μM) tubulin and GTP was 6 mM. The mixture was mixed, incubated on ice for 5 min, and microtubules were polymerized at 37°C for ∼1.5 hours. To stabilize the microtubules, polymerization buffer supplemented with taxol (MilliporeSigma) was added such that the final taxol concentration was ∼20 μM. The microtubules were pelleted on a benchtop centrifuge (Eppendorf) at 20,000 rcf for 8 min at room temperature. The supernatant was discarded and the pellet was resuspended in polymerization buffer with taxol by pipetting up and down with a cut tip. The microtubules were pelleted and resuspended again using an uncut tip. The concentration was approximated using Bradford reagent (Biorad) and diluted to ∼0.65 mg/mL (∼6 μM).

To assemble the dynein-dynactin-BICDR complex, the purified proteins were mixed in a 1:2:32 molar ratio (0.33 mg/mL dynein, 0.56 mg/mL dynactin, 0.97 mg/mL BICDR) in GF150 buffer supplemented with 1 mM DTT in 10 μL and incubated on ice for 15 min. To bind the complex to microtubules, 9 μL complex was mixed at room temperature with 5 μL microtubules and 9 μL binding buffer A (77 mM HEPES pH 7.2, 51 mM KCl, 13 mM MgSO_4_, 2.6 mM EGTA, 2.6 mM DTT, 7.68 mM AMPPNP, 13 μM taxol) such that the final concentrations of KCl and AMPPNP were 100 mM and 3 mM, respectively. After 5 min, the complex-bound microtubules were pelleted at 20,000 rcf for 8 min at room temperature. The pellet was resuspended in binding buffer B (30 mM HEPES pH 7.2, 60 mM KCl, 5 mM MgSO_4_, 1 mM EGTA, 1 mM DTT, 3 mM AMPPNP, 5 μM taxol, and 0.01% IGEPAL (MilliporeSigma)) using an uncut tip. 3.5 μL was applied to freshly glow-discharged Quantifoil R2/2 300-square-mesh gold grids (Quantifoil) in a Vitrobot IV (ThermoFisher) at 100% humidity and 20°C, incubated for 20 s, and blotted for 0.5-1.5 s before being plunged into liquid ethane.

### Cryo-EM

The samples were imaged using a FEI Titan Krios (300 kV) equipped with a K3 detector and energy filter (20 eV slit size) (Gatan) using automated data collection (ThermoFisher EPU) over 14 sessions. In total, 88,715 movies were acquired at 81,000 X magnification (1.11 Å/pixel, 100 μm objective aperture, ∼3 sec exposure, 53 frames, −1.2 to −3.6 μm defocus range), and 9,875 movies were acquired at 42,000 X magnification (2.13 Å/pixel, 70 objective μm aperture, ∼9 sec exposure, 53 frames, −1.5 to −4 μm defocus range). 1,846 of the low magnification movies were acquired at a 26° tilt of the stage. In all cases, the total fluence was ∼53 e^-^/Å^2^.

### Cryo-ET

Samples were prepared as for cryo-EM except that gold fiducials (BBI Solutions) pre-equilibrated in binding buffer B were added to the sample before freezing. Tomograms were acquired using a Glacios TEM (200 kV) equipped with a Falcon III detector (Thermo Fisher). SerialEM [Mastronarde, 2005] was used to acquire a tilt series between −60° and +60° in 3° increments at 57,000 X magnification (2.55 Å/pixel, 70 μm objective aperture). At each tilt angle, a movie containing 13 frames with an exposure of ∼3 e^-^/Å^2^/tilt. The total fluence for the tilt series was ∼120 e^-^/Å^2^ and the defocus range was −3 to −6 μm. Gain and motion correction was done using alignframes from IMOD [Kremer et al., 1996], with per-frame dose weighting. Tomogram alignment and reconstruction (back projection) was done using the ETomo interface of IMOD. The tomograms were subsequently binned by 4 and processed using a Wiener-like deconvolution filter as implemented in WARP [Tegunov and Cramer, 2019] (https://github.com/dtegunov/tom_deconv). The resulting tomograms were visualized in ChimeraX [Goddard et al., 2018] using the Hide Dust option set to ∼200. Complexes where all four motor domains were clearly defined were chosen for visualization. To color the density, the Color Zone option was used with the atomic models of dynein and dynactin, which were rigid body fit into the density and manually adjusted in cases where the motor domains were in different conformations.

### Microtubule subtraction and particle picking

Micrographs were generated from the cryo-EM movies after correcting for drift using MotionCorr2 [Zheng et al., 2017] implemented in Relion 3.1 [Zivanov et al., 2018], and the contrast transfer function (CTF) parameters were estimated using CTFFIND4 [Rohou and Grigorieff, 2015]. Microtubules were picked from the micrographs using the filament option in crYOLO [Wagner et al., 2020] with a model that was trained on manually picked micrographs (∼100). A different model was used for low- and high-magnification micrographs. The output coordinates along each microtubule were spaced by 81 Å. These coordinates were used in the Microtubule Relion-based Pipeline (MiRP) to reconstruct the microtubules [Cook et al., 2020]. Briefly, the coordinates were used to extract particles at 4x binning and a rolling average of the particles along a microtubule were generated to increase the contrast for subsequent classification steps. A model for 11 to 16 protofilament microtubules was used for supervised 3D classification in Relion 3.0, 3.1, or 4.0 [Zivanov et al., 2018; Kimanius et al., 2021]. The protofilament number (pf) distribution was 11 pf: 2.1% ± 0.3, 12 pf: 18.6% ± 3.2, 13 pf: 60.0% ± 1.6, 14 pf: 16.7% ± 1.7, 15 pf: 1.2% ± 0.2, 16 pf: 1.4% ± 0.3. After assigning the most likely pf number to each microtubule, the data was split and each type was processed independently. Sequential refinement of the helical parameters (Rot angle, X/Y shifts) following the MiRP protocol led to properly aligned and internally-consistent particles. The seam checking step was skipped since the misalignment of the seam was not detrimental to the quality of the subtraction described below. The data was re-extracted to generate raw particle stacks at 4x binning for local refinement in Relion 3.1 or 4.0, and then re-extracted without binning for a final local refinement.

Microtubules were subtracted using the *relion_particle_reposition* command-line interface in Relion 4.0 by providing the final alignment parameters and 3D density. Dynein-dynactin-BICDR particles were then picked from the subtracted micrographs using a crYOLO model that was pre-trained on ∼100 micrographs [Wagner et al., 2019]. Complexes that were too far from the microtubules were removed using the *extract_if_nearby* option in Starparser (https://github.com/sami-chaaban/starparser). Specifically, the closest microtubule coordinate was found for every complex coordinate, and those with distances that were greater than ∼550 Å were removed. From this point onwards, the data was processed in Relion 3.1 or Relion 4.0 unless otherwise stated.

### Processing

A processing pipeline can be found in Figure S2. Picked particles that were close to microtubules were extracted with binning such that the final pixel size was 3 Å/pixel. A reference was generated using a manually built model of the full dynein-dynactin-BICDR complex based on previous structural work [Urnavicius et al., 2018; Grotjahn et al., 2018], which was then filtered to 80 Å for 3D classification with alignment. Multiple classifications were performed in parallel, with each run varying in settings (ignore CTFs until first peak: Yes or No, mask diameter 780 Å or 820 Å, number of classes 6, 12, or 15, tau (T) value of 3 or 6). Particles from the best class in each run were pooled and duplicates were removed based on the rlnImageName column using the *remove_duplicates* option in Starparser. The number of particles remaining at this stage were 506,853 and 121,180 in the high-magnification and low-magnification set, respectively.

The particles were subjected to a global 3D refinement without a mask using a 60 Å filtered reference. A soft-edge mask was generated based on the resulting density and the refinement was allowed to proceed with the mask by choosing “Continue”. The high-magnification data were then re-extracted from the non-microtubule-subtracted micrographs at 1.17x binning (1.30 Å/pixel). Local refinement, Bayesian polishing, and a further local refinement yielded a consensus structure at 4.60 Å resolution. The dynein-tail/dynactin region was masked for local refinement and used for multiple signal subtractions to improve the local resolutions. In all cases, local refinement, defocus refinement, magnification refinement, beam-tilt refinement, and at least one round of 3D classification without alignment was performed to improve the resolution. The final maps were B-factor sharpened for model building (described below). EMDA was used to align the maps to the consensus dynein-tail/dynactin map [Warshamanage et al., 2021]. All masks, particle numbers, resolutions, classification parameters, and B-factors can be found in Figure S2.

To solve the structure of the motor domain, the signal for dynein-tail/dynactin was first subtracted by using the local refinement around dynactin described above as input to a signal subtraction with a mask around all four motor domains. In parallel, a local refinement of the motor domains using the same mask was performed, and the alignments from this run were imported into the signal subtracted particles using the *swap_columns* option in Starparser, by passing the following columns: rlnAnglePsi, rlnAngleRot, rlnAngleTilt, rlnNormCorrection, rlnLogLikeliContribution, rlnMaxValueProbDistribution, rlnNrOfSignificantSamples, rlnOriginXAngst, and rlnOriginYAngst. This provided a more accurate set of starting alignments for a subsequent 3D classification without alignment of the subtracted particles into two classes. For each class, which represented the staggered and aligned states, four parallel signal subtractions were performed on each motor domain. The particles were then joined from all eight signal subtraction jobs. A 3D refinement yielded a structure at 7.03 Å, which was further improved to 3.52 Å after defocus refinement, magnification refinement, beam-tilt refinement, and two rounds of 3D classification without alignment. To improve the density of the nucleotide in the AAA1 domain, further signal subtraction was performed with a mask around AAA1, AAA2, and AAA3. Local refinement and 3D classification without alignment resulted in improved density for AAA1. To improve the density of the stalk, the refinement of the motor domain was signal subtracted with a mask around the stalk and microtubule binding domain. 3D classification without alignment identified particles with the clearest density, which were reverted to the original box size for two rounds of local refinement and 3D classification. All masks, particle numbers, resolutions, classification parameters, and B-factors can be found in Figure S2.

For more challenging regions such as the BICDR-B and the LICs, the two magnification datasets were combined: the high-magnification data were re-extracted from microtubule-subtracted micrographs at ∼2x binning and combined with the low-magnification data (unbinned), for a shared pixel size of ∼2.13 Å/pixel. Local refinement yielded a consensus structure at 6.33 Å resolution. To better define the p150^Glued^ connecting density, a 3D classification (Tau fudge = 20, 15 classes) was performed and the class with the clearest signal (30,570 particles) was further locally refined (13.50 Å final resolution).

The consensus structure from the combined-magnification datasets was used for signal subtraction with a mask around dynactin or the BICDR-B N-terminus. The dynactin-focused particles were used for cryoDRGN classification of the LICs (described below) and for further signal subtraction with a mask around the pointed end. A final round of signal subtraction around the density for BICDR-B was performed in order to isolate particles with the clearest density, before reverting to the pointed-end signal subtraction. This resulted in an improved density for BICDR-B at the pointed end. For the BICDR-B N-terminus, the density was improved after signal subtraction with defocus refinement, magnification refinement, beam-tilt refinement, and 3D classification without alignment. All masks, particle numbers, resolutions, classification parameters, and B-factors can be found in Figure S2.

To better define the motor-domain interactions, the consensus refinement from the combined-magnification datasets was signal subtracted with a mask around all four motor domains. In parallel, a local refinement of the motor domains using the same mask was performed, and the alignments were imported into the signal subtracted particles using the *swap_columns* option in Starparser as described above. A 3D classification separated particles from the staggered and aligned states. For each state, a mask around dynein-A1/A2 and dynein-B1/B2 was used for signal subtraction, followed by 3D classification without alignment and 3D refinement to improve the density. All masks, particle numbers, resolutions, classification parameters, and B-factors can be found in Figure S2.

All structures were post-processed in Relion by including the MTF of the detector. The maps were also scaled to the correct pixel size, which was calibrated by comparing the density for a tubulin dimer to a previous structure of taxol-bound tubulin [Kellogg et al., 2017]. Specifically, the maps were fit to each other using the Fit in map option in Chimera, and the pixel size scaling with the highest correlation was used. The final pixel size for the high magnification and combined-magnification datasets were 1.24 Å/pixel (originally 1.30 Å/pixel) and 2.04 Å/pixel (originally 2.13 Å/pixel), respectively. This scaling also resulted in a match to previous dynactin structures [Lau et al., 2021].

The composite map in Figure 1D was built by summing the maps in Chimera [Pettersen et al., 2004]. For the motor domains, the map with improved density for the stalk was fit into each of the four positions in the motor domain-pair maps. Although the raw data shows that there’s flexibility in the orientation of the complex relative to the microtubule, for illustrative purposes the composite map was overlaid onto the microtubule to match the configuration of the microtubule binding domains with the tubulin dimers [Redwine et al., 2012; Lacey et al., 2019]. All colours were rendered in ChimeraX.

### Model building and refinement

Building was performed in COOT [Emsley et al., 2010] and refinement was done in PHENIX [Afonine et al., 2018]. All structure predictions were performed using Alphafold through a local installation of Colabfold 1.2.0 [Mirdita et al., 2021], running MMseqs2 [Mirdita et al., 2019] for homology searches and Alphafold2 [Jumper et al., 2021] or Alphafold2-Multimer [Evans et al., 2021] for the predictions of single or multiple chains, respectively. When used to guide model building, predictions were generated with amber relaxation [Hornak et al., 2006]. All refinement statistics can be found in Table S1 and Table S2.

A starting model of the dynein motor domain was generated by using a previous structure of dynein in the phi conformation (PDB: 5NUG) [Zhang et al., 2017] and manually adjusting it to match the crystal structure of S. cerevisiae dynein with AMPPNP (PDB: 4W8F) [Bhabha et al., 2014] in COOT. The model was then rebuilt into the density and challenging regions were guided by Alphafold predictions of the individual AAA+ subdomains. Nucleotides were placed in the AAA+ domains as follows (AAA1: ADP, AAA2: ATP, AAA3: AMPPNP: AAA4: AMPPNP). The model was then manually inspected before refinement.

The dynein-A2/B1 interaction in the staggered state was built by placing into the density the refined motor domain from above with an Alphafold prediction of the LIC (O43237) Ras-like domain, before refinement in COOT.

The N-terminus of the dynein tails were built using previous structures of dynactin/dynein-tails as a reference (PDB: 6F1T) [Urnavicius et al., 2018] and refined. An Alphafold prediction of the dynein intermediate chain (Q13409) WD40 domain was built into the structure and refined.

Dynactin subunits were built and refined using previous models as a reference (PDB: 6ZNL, 6F1T) [Lau et al., 2021; Urnavicius et al., 2018], as well as Alphafold predictions to guide challenging regions, and were manually inspected and refined. The pointed end complex includes actin-related protein 11 (Arp11, ACTR10), p62 (DCTN4), p25 (DCTN5), and p27 (and DCTN6), as well as an N-terminal fragment of p50 (DCTN2) [Lau et al., 2021]. The Arp1/actin filament includes eight copies of actin-related protein 1 (ARP1, also known as ACTR1A) and one copy of *β*-actin [Urnavicius et al., 2018]. The barbed end includes CAPZ*α* and CAPZ*β* [Urnavicius et al., 2018].

The structure of full-length BICDR was first predicted in Alphafold on two copies of BICDR (BICDL1: A0JNT9), and was flexibly fit into a low-resolution map of BICDR-A in COOT using the registry defined by the position of Trp166. The N-terminus containing the CC1 and CC2 box was then built into high-resolution density and refined. The C-terminus containing the Spindly motif was guided by a prediction of the pointed end complex (described below). The BICDR-B model was built by flexible fitting of the BICDR-A model into a low-resolution map of BICDR-B after adjusting the registry based on the position of Trp166.

The LIC C-terminal alpha helices were built from a prediction of two copies of the BICDR N-terminus (BICDL1: A0JNT9) and two copies of the LIC (O43237) C-terminus, before being refined in COOT.

To generate the composite structure, a copy of the motor domain was placed in each monomer position in the composite map. The C-terminus of the dynein tails were built by first generating predictions of the dynein heavy chain (Q14204) helical bundles and placing them into the density, before stitching them together. The model was then refined in COOT into the density of each tail and stitched to the N-termini of the dynein tails and the motor domains. The dimerization domain was built by generating a prediction of two copies of the N-terminus of dynein, which was refined into the density in COOT. The complex of dynein light chain ROBL1 (DYNLRB1) and the dynein intermediate chain was flexibly fit into the density from a previous structure (PDB: 6F1T) [Urnavicius et al., 2018]. The shoulder was rigid body docked from a previous structure (PDB: 6ZNL) [Lau et al., 2021], which includes two copies of p150^Glued^, four copies of p50 (DCTN2), and two copies of p24 (DCTN3). The stalk of the motor domains were built by flexible fitting of the *T. thermophila* axonemal dynein stalk (PDB: 7K58) [Rao et al., 2021] before mutating the residues to match the human sequence. The stalk was then stitched to the structure of the human dynein microtubule binding domain (PDB: 6RZB) [Lacey et al., 2019]. The stalk and microtubule binding domain were refined into the density of each motor position before being stitched to the motor domains.

### CryoDRGN classification

To help reveal densities for the flexible LIC loops, the consensus structure from the combined-magnification datasets that was signal subtracted around dynactin was used as input for multiple rounds of cryoDRGN analysis [Zhong et al., 2021]. First, the particles were binned from 400 pixels to 128 pixels (6.75 Å/pixel final pixel size) and used for training an 8-dimensional latent variable model with 3 hidden layers and 256 nodes in the encoder and decoder networks. The latent space was visualized with the *analyze* function on epoch 50 and 3 clusters were extracted using a gaussian mixture model (GMM). The particles from the cluster that yielded the best 3D refinement were moved forward for two more rounds of training, GMM clustering, and 3D refinement, but with binning to 256 pixels (3.33 Å/pixel final pixel size) and training with 3 hidden layers and 1024 nodes in the networks. Finally, cryoDRGN *landscape* analysis was performed on the latent space using a custom mask in the region encompassing the LICs, with 20 clusters extracted using ward linkage. The mean volume for each cluster was manually inspected for connecting density between the LIC Ras-like domain and the BICDR N-termini.

Previous data (dynein-tail/dynactin/BICDR and dynein-tail/dynactin/Hook3) [Urnavicius et al., 2018] was reanalyzed by binning the consensus particles from 432 pixels to 256 pixels (2.26 Å/pixel final pixel size) and training an 8-dimensional latent variable cryoDRGN model with 3 hidden layers and 1024 nodes in the encoder and decoder networks. The latent space was visualized with the *analyze* function on epoch 50 and epoch 41 in the BICDR and Hook3 datasets, respectively, and 20 clusters were extracted using k-means analysis. The structure representing the center of each cluster was manually inspected for the presence of a second adaptor.

### Motor domain model alignments

To analyse the nucleotide pockets in AAA1 and AAA3, the large subdomains were aligned to the crystal structure of *S. cerevisiae* dynein AMPPNP [Bhabha et al., 2014] and *D. discoidium* dynein ADP [Kon et al., 2012] using matchmaker in Chimera. The difference between our structure and the crystal structure was visualized by drawing arrows between the coordinates of corresponding residues using a custom Python script. To determine which pair of residues the arrows should be drawn between, the sequence of the subdomains of interest were aligned using Clustal Omega [Sievers et al., 2011]. Pairs that were more than 12 Å apart were ignored, which likely represent residues that were poorly aligned.

### Mass photometry

To analyse the oligomerization of BICDRs in solution, we subjected the purified protein to mass photometry using the Refeyn OneMP instrument (Refeyn Ltd.). BICDR1 was diluted to 75 nM in GF150 buffer and 10 μL was imaged. The movies were processed using DiscoverMP and the mass was estimated by fitting a Gaussian distribution to the data.

### Cargo adaptor structure prediction

The pointed end-BICDR complex was predicted using Colabfold 1.2.0 (10 recycles, no templates) with the following sequences: DCTN4 (Uniprot ID Q9UJW0), DCTN6 (O00399), DCTN5 (Q9BTE1), ACTR10 (I3LHK5), and two copies of BICDR (BICDL1; A0JNT9) fragment 205-394. The predicted align error (PAE) with respect to Leu347 was mapped onto the predicted structure using a custom Python script that extracts the values from the plot and generates an attribute file for ChimeraX visualization.

Cargo adaptors were predicted by running Colabfold 1.2.0 (no templates) on two copies each of the following sequences: BICDR1 (BICDL1: A0JNT9), BICD2 1-423 (Q8TD16) Spindly (SPDL1: Q96EA4), TRAK1 1-450 (Q9UPV9), Hook3 (Q86VS8), RAB11FIP3 (RFIP3: O75154), Cracr2a (EFC4B: Q9BSW2), JIP3 1-700 (Q9UPT6), RILP (Q96NA2). For visualization, the models were manually linearized such that the coiled-coils run parallel to each other. Specifically, the predictions were broken up into fragments at flexible loops, rotated to be parallel, and stitched again in COOT. The PAEs and predicted local distance difference tests (pLDDTs) at the coiled-coil breaks preceding the Spindly motif can be found in Figure S10. The CC1 box, CC2 box, and Spindly motif were manually annotated in Figure S8C based on the presence of the motifs in the appropriate regions of the coiled-coils [Sacristan et al., 2018].

The Hook3 CC2 box region was predicted by running Colabfold 1.2.0 (no templates) on two copies of Hook3 172-287 (Q86VS8), one copy of dynein heavy chain 576-864 (Q14204), and one copy of dynein intermediate chain (Q13409) 226-583. The PAE relative to Hook3 His200 was mapped onto the prediction as above for the pointed end complex.

## Acknowledgements

We thank S. Scheres for help with micrograph-level signal subtraction in Relion; C.K. Lau for helpful discussions; the MRC Laboratory of Molecular Biology Electron Microscopy Facility for access and support of electron microscopy sample preparation and data collection; J. Grimmett and T. Darling for providing scientific computing resources; H.E. Foster and C. Ventura Santos for help with cryo-ET; F. Abid Ali, K. Singh, and C.K. Lau for critical reading of the manuscript; Henriques, R. for the Biorxiv template. This work was supported by Wellcome [210711/Z/18/Z], the Medical Research Council, as part of United Kingdom Research and Innovation (also known as UK Research and Innovation) [MRC file reference number MC_UP_A025_1011], and the EMBO Postdoctoral Fellowship [ALTF 334-2020] to S.C. For the purpose of open access, the author has applied a CC BY public copyright license to any Author Accepted Manuscript version arising.

## Author Contributions

S.C. performed experiments/analysis and prepared figures. S.C. and A.P.C. conceived the project and wrote the manuscript.

## Code Availability

Custom scripts, including the Starparser package, are available at https://github.com/sami-chaaban.

## Data Availability

Atomic coordinates and cryo-EM maps have been deposited in the Protein Data Bank (PDB) / Electron Microscopy Data Bank (EMDB) under accession codes 7Z8F/14549 (composite dynein-dynactin-BICDR structure), 7Z8G/14550 (dynein motor domain), 7Z8H/14551 (dynein AAA1-3), 7Z8I/14552 (Barbed end/BICDR-A), 7Z8J/14553 (BICDR-A/dynein-A2), 7Z8K/14555 (BICDR-B/dynein-A1), 7Z8L/14556 (dynein motor domain/LIC), and 7Z8M/14559 (pointed end/BICDR-A).

---

**Table S1.** Cryo-EM data collection, refinement, and validation statistics of the dynactin/dynein-tail regions.

| Region | Pointed end/<br>Spindly motif | BICDR-A<br>Dynein-A2 Tail | Barbed end/<br>BICDR-A CC1 box | BICDR-B/<br>Dynein-A1 Tail |
| --- | --- | --- | --- | --- |
| EMDB | 14559 | 14553 | 14552 | 14555 |
| PDB | 7Z8M | 7Z8J | 7Z8I | 7Z8K |
| **Data collection/processing** |  |  |  |  |
| Magnification | 81000 | 81000 | 81000 | 81000 |
| Voltage (kV) | 300 | 300 | 300 | 300 |
| Electron exposure (e<sup>-</sup>/Å<sup>2</sup>) | 53 | 53 | 53 | 53 |
| Defocus range (μm) | 1.2 - 3.6 | 1.2 - 3.6 | 1.2 - 3.6 | 1.2 - 3.6 |
| Pixel size (Å) | 1.2445 | 1.2445 | 1.2445 | 1.2445 |
| Symmetry imposed | C1 | C1 | C1 | C1 |
| Initial particle images (no.) | 506853 | 506853 | 506853 | 506853 |
| Final particle images (no.) | 165019 | 78061 | 179660 | 39644 |
| Map resolution (Å) | 3.37 | 3.93 | 3.30 | 4.37 |
| FSC threshold | 0.143 | 0.143 | 0.143 | 0.143 |
| **Refinement** |  |  |  |  |
| Map-model FSC (Å) |  |  |  |  |
| Masked/unmasked | 3.47/3.55 | 3.91/3.95 | 3.62/3.56 | 7.58/8.50 |
| FSC threshold | 0.5 | 0.5 | 0.5 | 0.5 |
| Map sharpening B factor (Å<sup>2</sup>) | -55 | -100 | -55 | -67 |
| Model composition |  |  |  |  |
| Non-hydrogen atoms | 17689 | 20293 | 43524 | 14477 |
| Protein residues | 2273 | 2487 | 5372 | 2918 |
| Ligands | ADP:3, ATP:1, ZN: 3 | - | ADP:6 | - |
| B factors (Å<sup>2</sup>) |  |  |  |  |
| Protein (min/max/mean) | 13.53/180.03/75.70 | 6.17/181.83/75.39 | 0.00/228.11/60.21 | 0.00/986.10/158.63 |
| Ligands (min/max/mean) | 10.20/126.82/63.73 | - | 20.79/75.00/44.34 | - |
| R.m.s. deviations |  |  |  |  |
| Bond lengths (Å) (#>4σ) | 0.005 (0) | 0.006 (2) | 0.005 (1) | 0.005 (1) |
| Bond angles (°) (#>4σ) | 1.134 (0) | 1.118 (2) | 1.055 (1) | 1.109 (0) |
| Validation |  |  |  |  |
| MolProbity score | 1.37 | 1.22 | 1.12 | 0.74 |
| Clashscore | 3.01 | 3.45 | 2.37 | 0.77 |
| Poor rotamers (%) | 0.05 | 0.18 | 0.15 | 0.00 |
| Ramachandran plot |  |  |  |  |
| Favored (%) | 95.91 | 97.63 | 97.52 | 98.06 |
| Allowed (%) | 4.09 | 2.37 | 2.48 | 1.94 |
| Disallowed (%) | 0.00 | 0.00 | 0.00 | 0.00 |

**Table S2.** Cryo-EM data collection, refinement, and validation statistics for the dynein motor domain regions.

| Region | Dynein<br>Motor domain | Dynein<br>AAA1-2-3 | Dynein-A2/B1 |
| --- | --- | --- | --- |
| EMDB | 14550 | 14551 | 14556 |
| PDB | 7Z8G | 7Z8H | 7Z8L |
| **Data collection/processing** |  |  |  |
| Magnification | 81000 | 81000 | 81000, 42000 |
| Voltage (kV) | 300 | 300 | 300 |
| Electron exposure (e<sup>-</sup>/Å<sup>2</sup>) | 53 | 53 | 53 |
| Defocus range (μm) | 1.2 - 3.6 | 1.2 - 3.6 | 1.2 - 4.0 |
| Pixel size (Å) | 1.2445 | 1.2445 | 2.0378 |
| Symmetry imposed | C1 | C1 | C1 |
| Initial particle images (no.) | 506853 | 506853 | 628033 |
| Final particle images (no.) | 351879 | 53424 | 105050 |
| Map resolution (Å) | 3.52 | 3.41 | 4.90 |
| FSC threshold | 0.143 | 0.143 | 0.143 |
| **Refinement** |  |  |  |
| Map-model FSC (Å) |  |  |  |
| Masked/unmasked | 3.65/3.82 | 3.55/3.60 | 6.43/6.99 |
| FSC threshold | 0.5 | 0.5 | 0.5 |
| Map sharpening B factor (Å<sup>2</sup>) | -70 | -75 | -112 |
| Model composition |  |  |  |
| Non-hydrogen atoms | 24656 | 8811 | 16513 |
| Protein residues | 3047 | 1083 | 3311 |
| Ligands | Mg:1, AMPPNP:2,<br>ADP:1, ATP:1 | Mg:1, AMPPNP:1,<br>ADP:1, ATP:1 | Mg:1, AMPPNP:2,<br>ADP:1, ATP:1 |
| B factors (Å<sup>2</sup>) |  |  |  |
| Protein (min/max/mean) | 35.76/152.59/63.47 | 8.85/113.21/47.93 | 35.76/152.59/65.90 |
| Ligands (min/max/mean) | 47.45/57.15/51.63 | 8.41/57.20/36.98 | 47.45/57.15/51.63 |
| R.m.s. deviations |  |  |  |
| Bond lengths (Å) (#>4σ) | 0.005 (0) | 0.005 (0) | 0.012 (2) |
| Bond angles (°) (#>4σ) | 1.073 (1) | 1.128 (0) | 1.536 (3) |
| Validation |  |  |  |
| MolProbity score | 1.20 | 1.29 | 0.61 |
| Clashscore | 3.41 | 3.71 | 0.29 |
| Poor rotamers (%) | 0.18 | 0.10 | 0.00 |
| Ramachandran plot |  |  |  |
| Favored (%) | 97.70 | 97.31 | 98.79 |
| Allowed (%) | 2.30 | 2.69 | 1.21 |
| Disallowed (%) | 0.00 | 0.00 | 0.00 |

**Figure S1.**
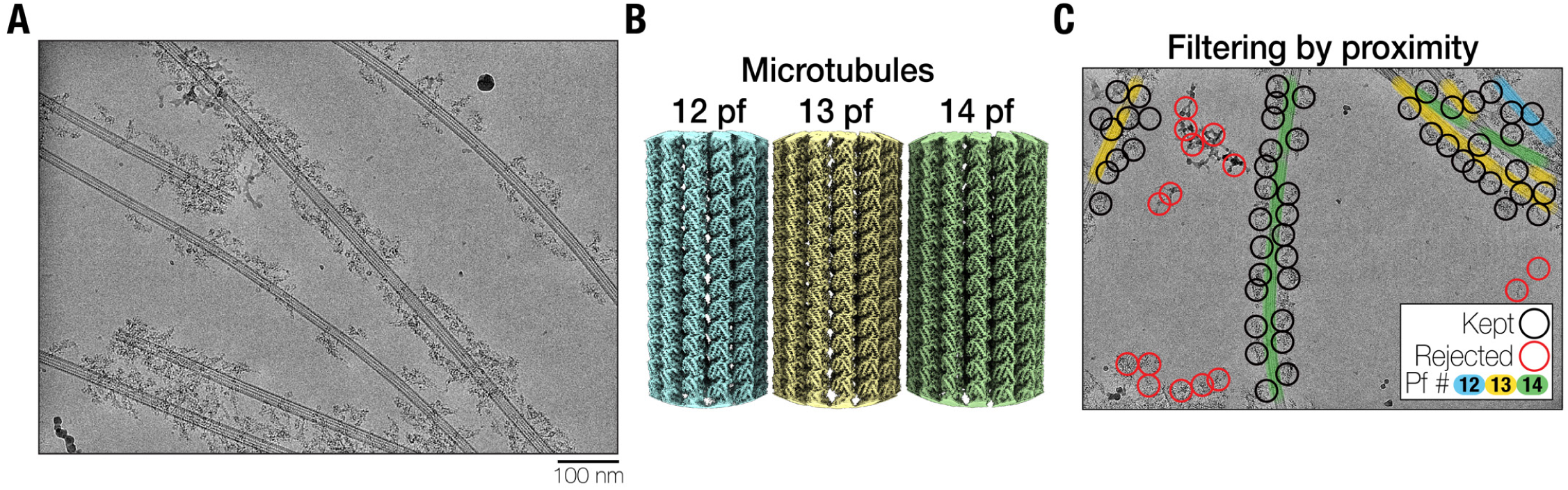
Cryo-EM processing of dynein-dynactin-BICDR micrographs. (A) An example micrograph. A pseudo-flat-field correction has been applied to normalize the intensity across the micrograph for visualization purposes (i.e. dividing the image by its gaussian blur). (B) The density maps of the 12, 13, and 14 protofilament (pf) microtubules. Not shown are the 11, 15, and 16 pf microtubules. (C) An annotated micrograph showing picked particles that were kept (black) or rejected (red) based on their proximity to the microtubules (blue, yellow, and green for 12, 13, and 14 pfs, respectively).

**Figure S2.**
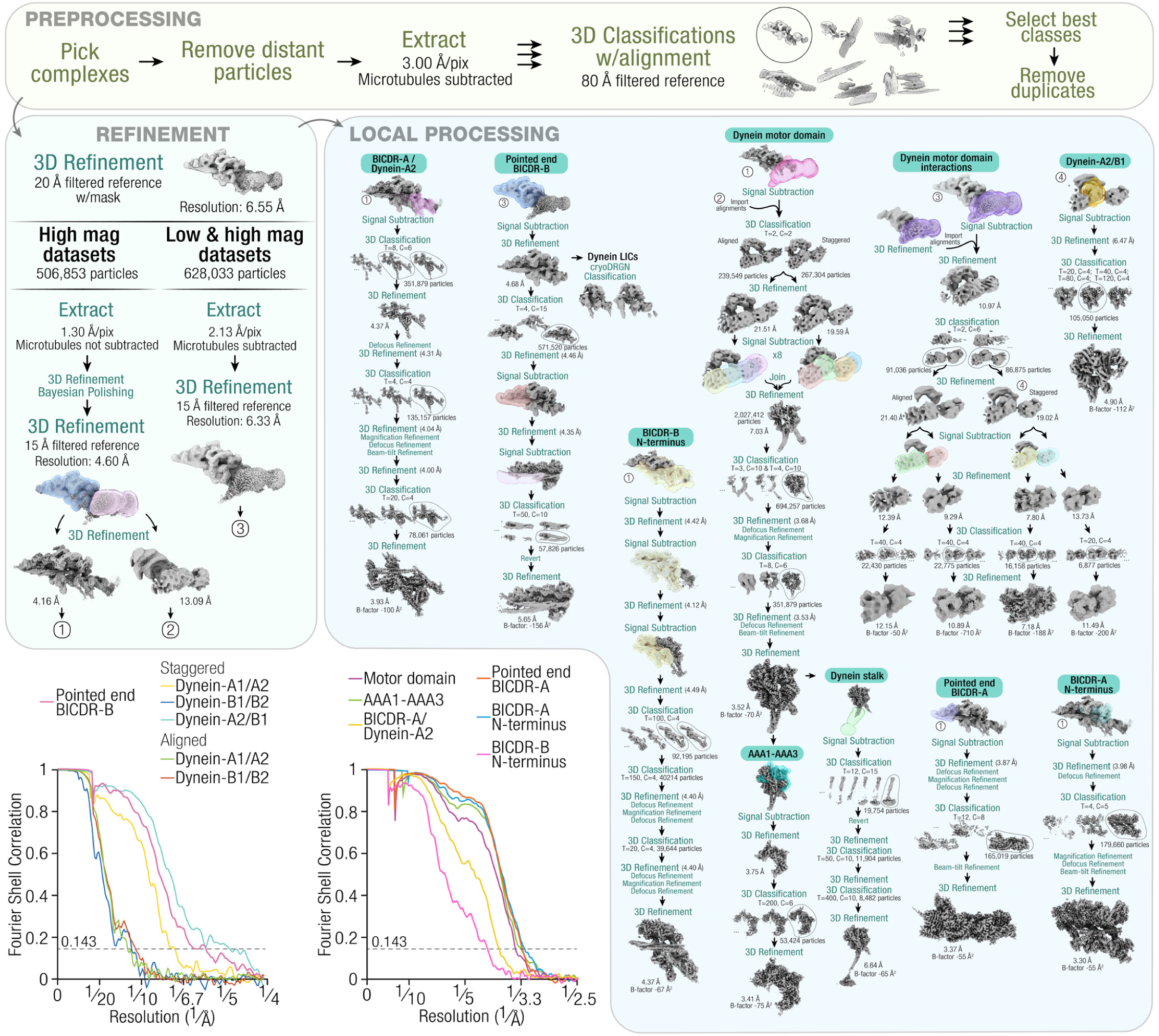
The processing pipeline for single particle analysis of the dynein-dynactin-BICDR complex. (T = tau fudge, C = number of classes). 3D classifications are without alignment unless otherwise specified. All defocus, magnification, and beam-tilt refinements were immediately followed by a 3D refinement (not shown). Plots show the gold standard Fourier shell correlation. The dotted horizontal line shows the 0.143 cutoff.

**Figure S3.**
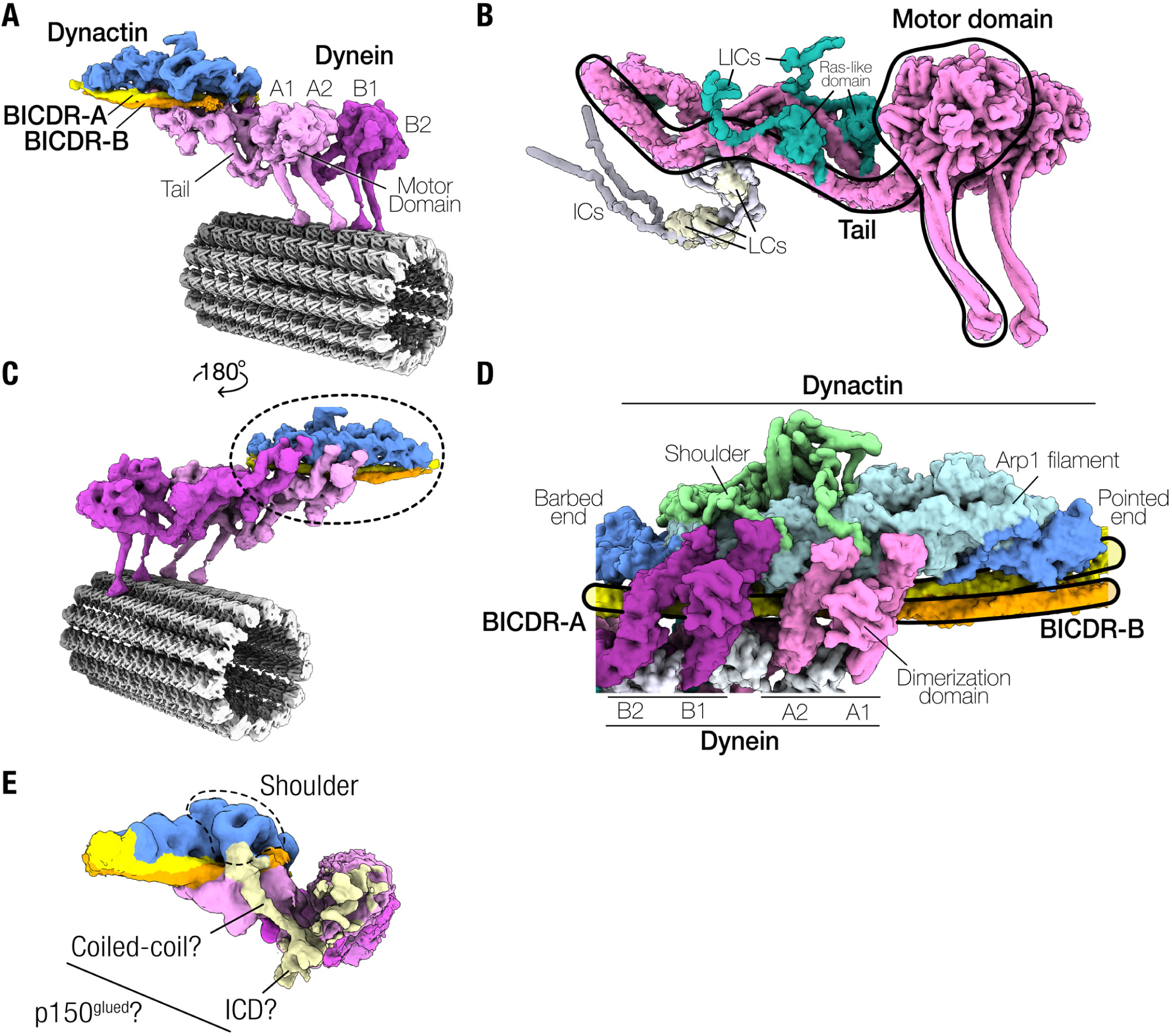
Overview of the dynein-dynactin-BICDR complex. (A) The composite density map of dynein-dynactin-BICDR overlaid on the reconstructed microtubule, showing the position of individual dynein motor domains (dynein-A1/2, dynein-B1/2), with their tails extending towards dynactin. (B) A molecular surface representation of a dynein dimer, showing the binding site the accessory proteins, including the LIC, which docks in the middle of the tail at the Ras-like domain and extends flexible loops. Note that the extended N-terminal domains of the intermediate chains (ICs), the associated light chains (LCs), and the flexible loops from the LICs are unresolved in the structure. (C) The composite density map of the complex shown from behind, where dynein’s tails can be seen binding in the grooves of dynactin’s Arp1 filament, which are spanned by the adaptors BICDR-A and BICDR-B. (D) A molecular surface representation of the back side of dynactin, showing the position of dynactin’s Arp1 filament and barbed/pointed ends relative to the dynein tails and BICDRs. (E) A 3D classification result on the whole complex from the combined magnification dataset showing density connecting the shoulder domain to the globular density near dynein-A1, which may represent the Inter-Coiled Domain (ICD) of p150^Glued^.

**Figure S4.**
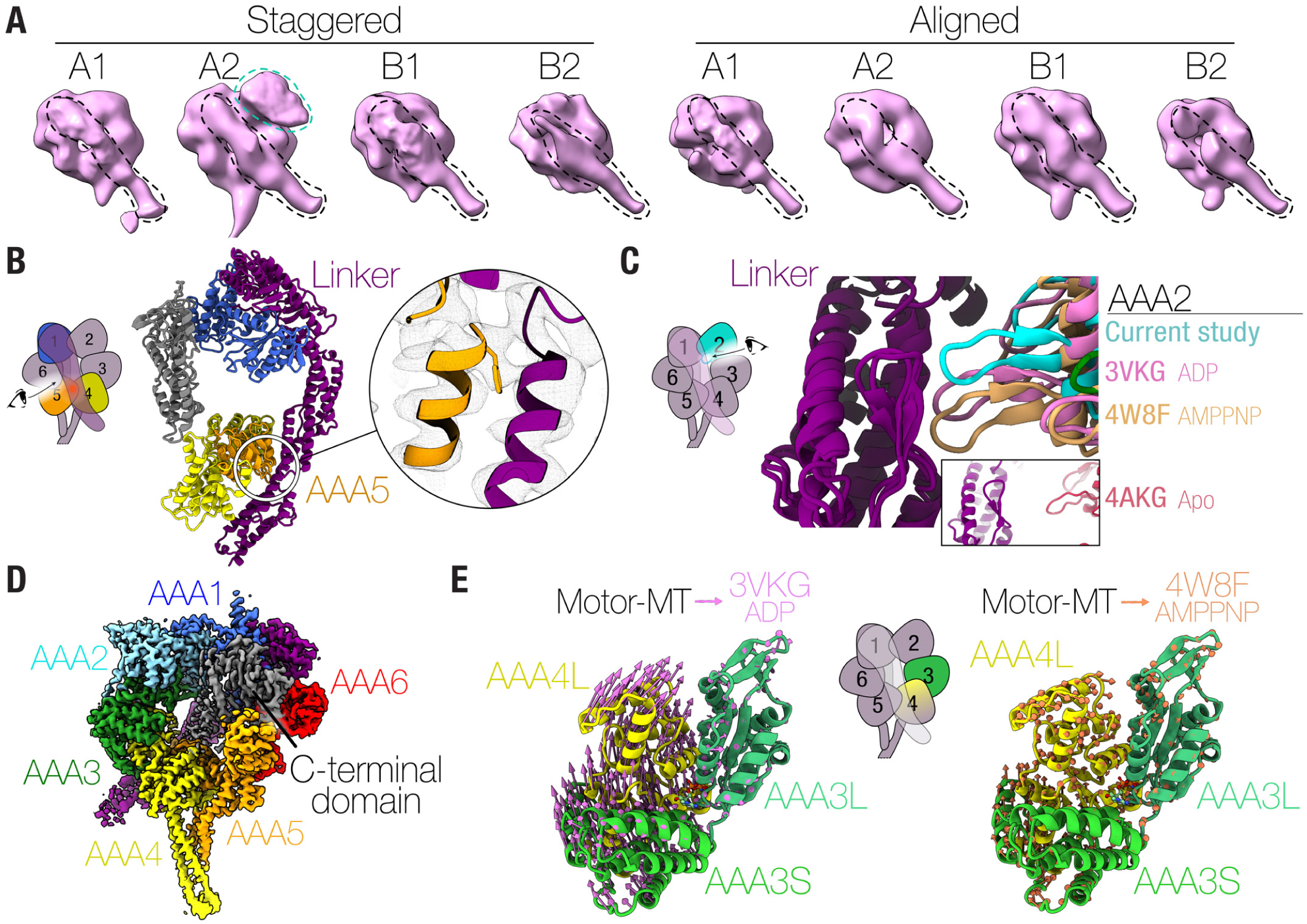
Conformation of the dynein motor domain. (A) A side view of the four motor domains from both major conformations (staggered and aligned) displaying a straight linker (dotted black line). Note the extra density on dynein-A2 in the staggered state, which represents the LIC of dynein-B1 (dotted teal line). (B) A front view of the motor domain only showing the linker (purple), AAA1 (blue), AAA4 (yellow), AAA5 (orange) and the C-terminal domain (gray). The inset shows the conserved Phe3629 in AAA5 binding the linker [Schmidt et al., 2012]. (C) A close-up view of the linker-AAA2 interaction overlaid with the crystal structures of *D. discoideum* dynein-ADP (PDB: 3VKG) [Kon et al., 2012] and *S. cerevisiae* dynein-AMPPNP (PDB: 4W8F) [Bhabha et al., 2014] aligned at the linker. The inset shows the crystal structure of *S. cerevisiae* dynein-Apo (PDB: 4AKG) [Schmidt et al., 2012], which lacks nucleotides in AAA1 and AAA3 and the linker is undocked from AAA2. (D) The density map view of the opposite face from the linker of dynein, where the C-terminal domain lies. (E) The domain movements in the nucleotide pocket of AAA3 represented by arrows after alignment of our structure (Motor-MT) to the crystal structure of *D. discoideum* dynein-ADP (left; PDB: 3VKG) [Kon et al., 2012], and *S. cerevisiae* dynein-AMPPNP (right; PDB: 4W8F) [Bhabha et al., 2014] aligned at AAA3L.

**Figure S5.**
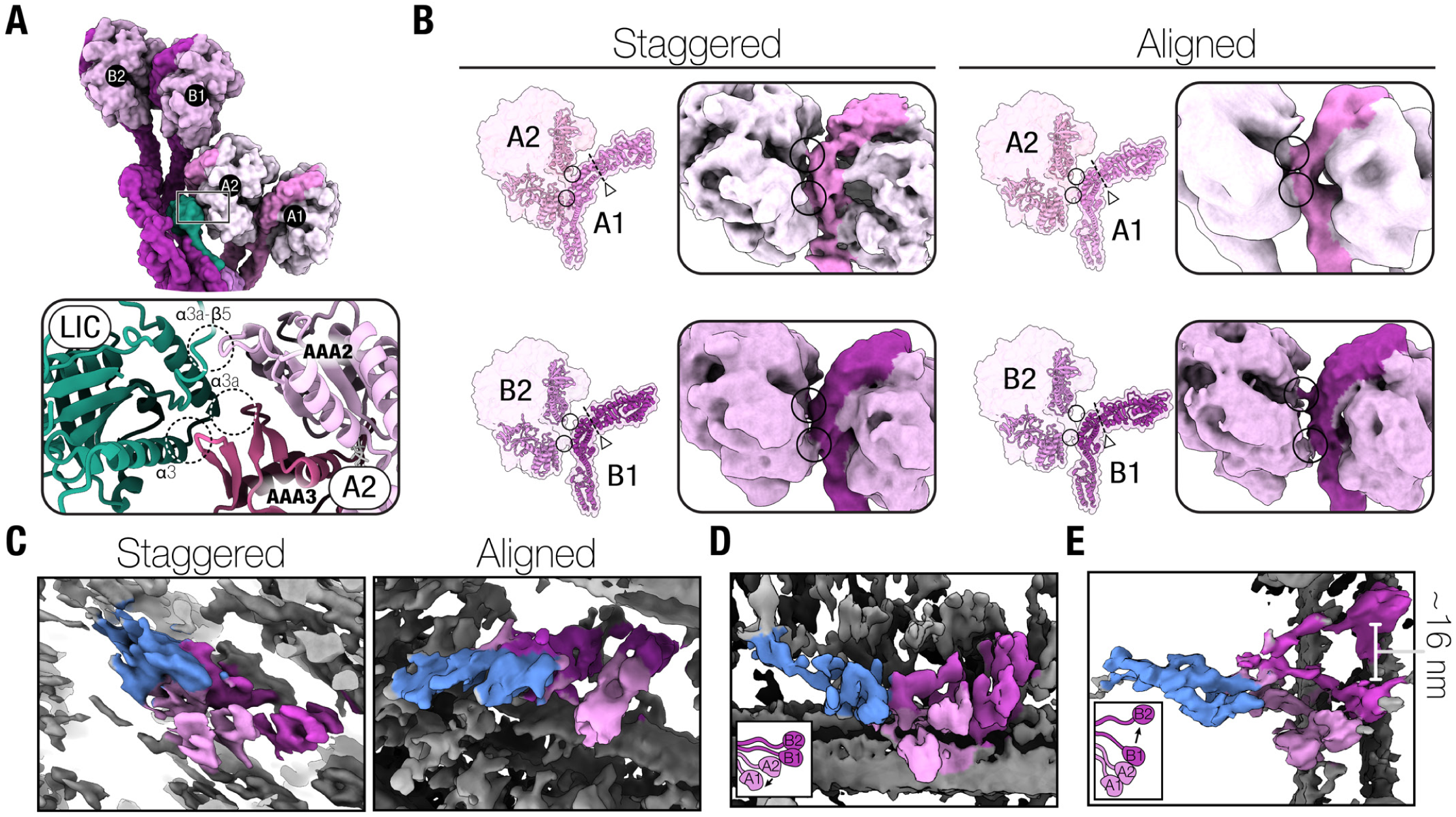
Dynein heavy chain interactions and conformational heterogeneity. (A) A close-up view of the interaction between dynein-A2 and the LIC of dynein-B1, highlighting the three potential interacting sites on the LIC. (B) The interaction between dynein motor domains in the staggered (left) and aligned (right) states is shown as a molecular surface representation and 3D density, where the linker is a darker colour in the latter. The triangles point to the hinge in the linker (dotted line). (C) Tomograms of dynein-dynactin-BICDR complexes on microtubules focused on individual complexes. The dynein and dynactin densities have been coloured pink/purple and blue, respectively. (D) Complexes in the tomograms that deviate from the staggered and aligned states. (E) An example tomogram where the complex has a large separation between the motor domains of dynein-B1 and B2.

**Figure S6.**
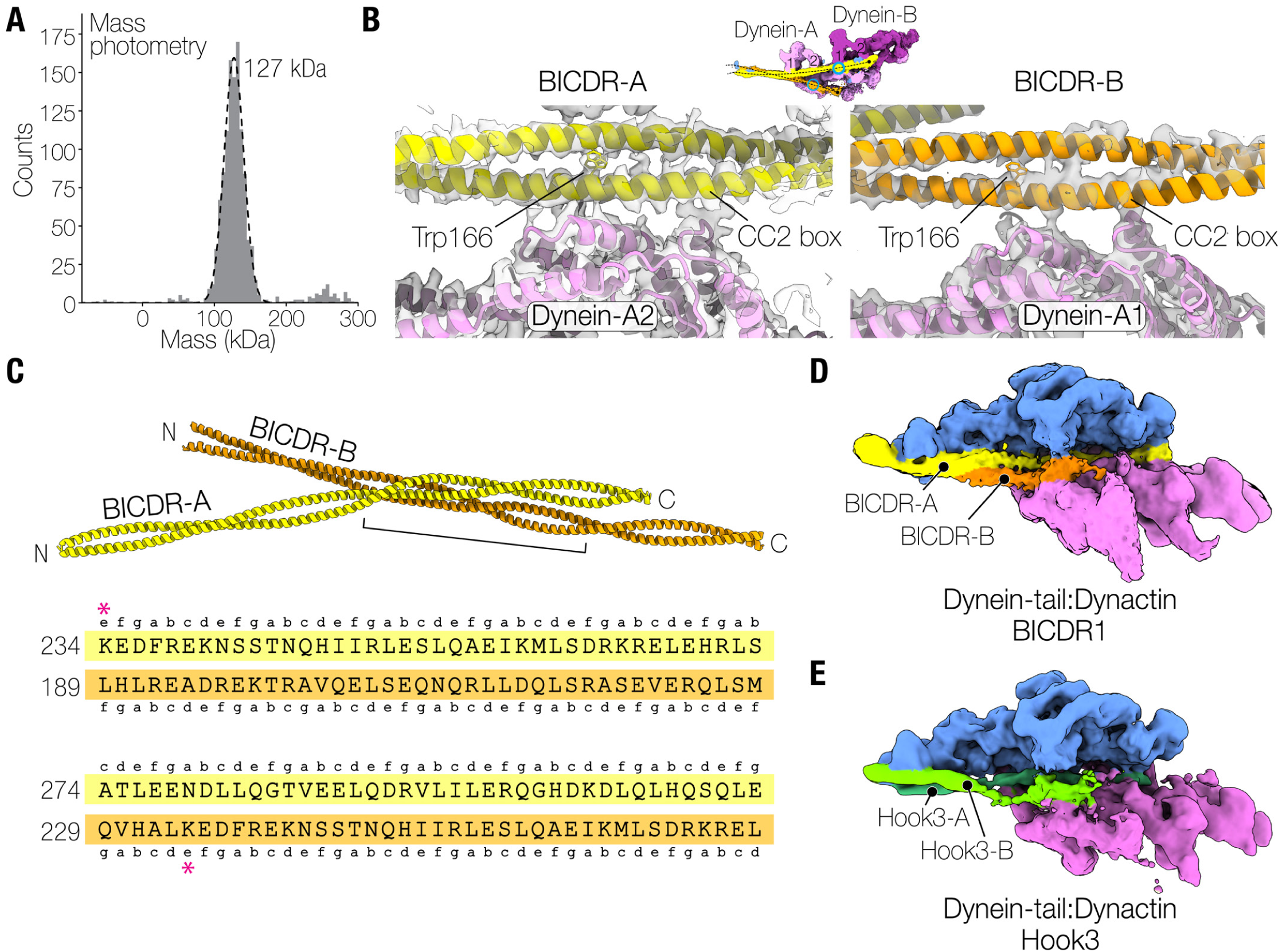
Cargo adaptor arrangements and interactions. (A) Mass photometry result of BICDR on its own, showing a major peak at 127 kDa (expected molecular weight is 130 kDa) and a minor peak at 260 kDa. (B) The density maps of BICDR-A and BICDR-B showing the bulky density at Trp166 and the location of the CC2 box near dynein-A2 and dynein-A1, respectively. (C) The interaction interface of the two BICDRs is shown based on their registry in the structure. Lowercase letters refer to the position in the heptad repeat of the coiled-coil, predicted with LOGICOIL [Vincent et al., 2013]. The red asterisk at K234 shows the offset between the two BICDRs. (D) A 3D classification of dynein-tail-dynactin-BICDR [Urnavicius et al., 2018] which shows density for a second adaptor. (E) A 3D classification of dynein-tail-dynactin-Hook3 [Urnavicius et al., 2018] showing that the second coiled-coil belongs to a second adaptor.

**Figure S7.**
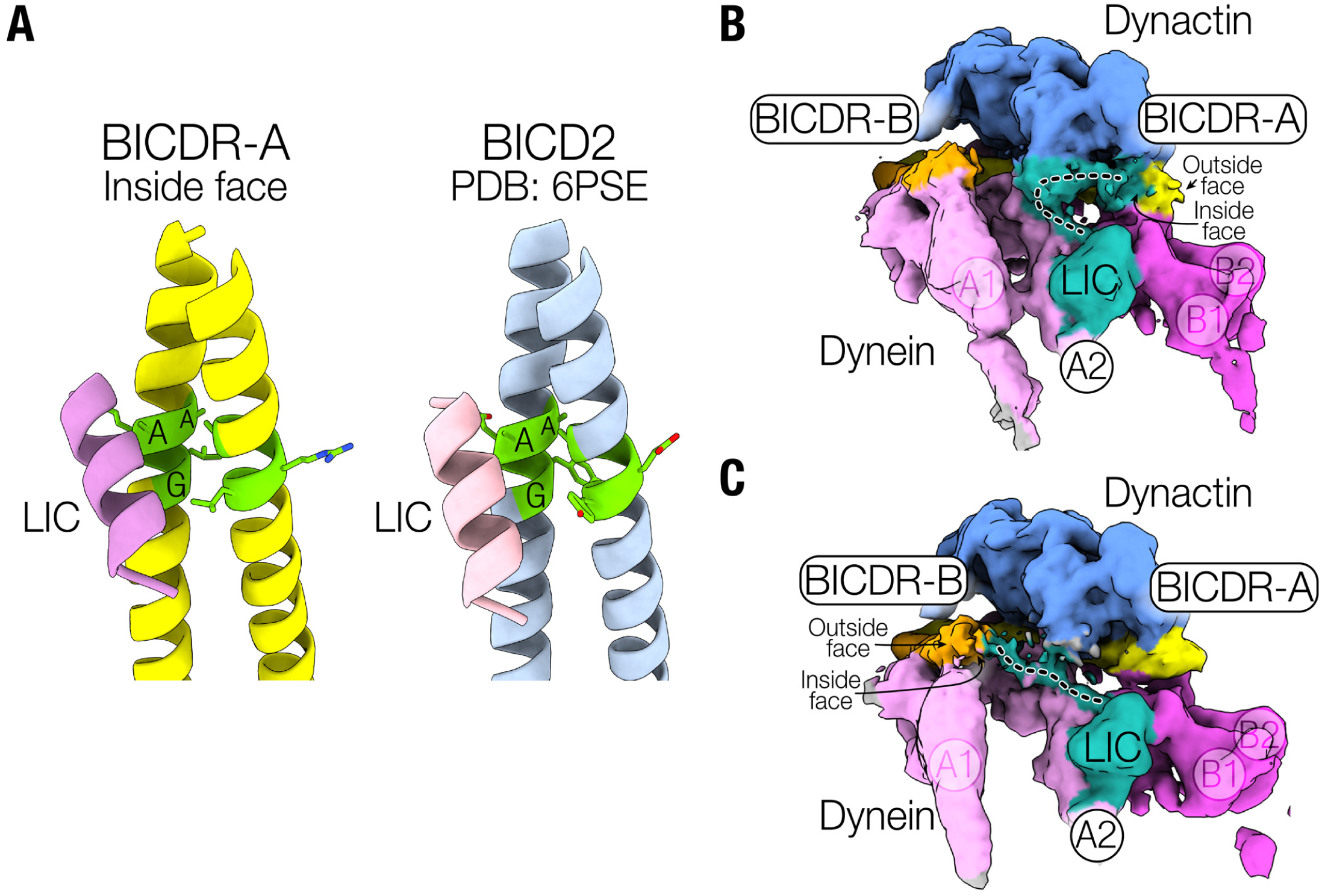
CC1 box interactions. (A) The LIC helix after fitting to the density on the inside face of the BICDR-A CC1 box relative to the registry of BICDR in our structure (left), highlighting the conserved Ala116, Ala117, and Gly120 of the AAXXG motif. On the right is a similar view of the BICD2-LIC crystal structure (PDB: 6PSE) [Lee et al., 2018]. (B) 3D classes showing the LIC of dynein-A2 connecting to BICDR-A (top) and BICDR-B (bottom).

**Figure S8.**
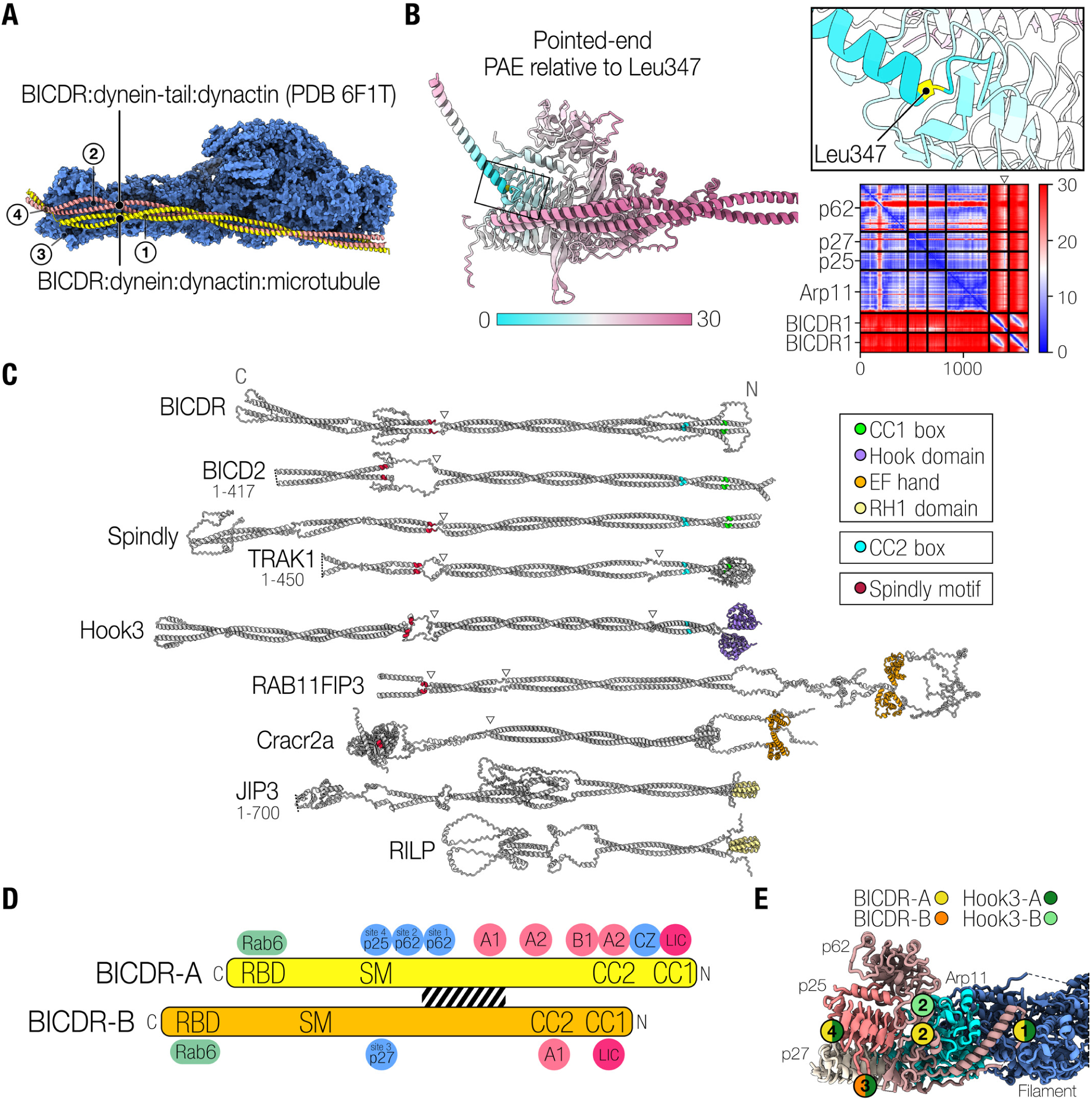
Pointed end interactions. (A) An overlay of our structure with the previous dynein-tail-dynactin-BICDR structure (PDB: 6F1T) [Urnavicius et al., 2018], aligned at dynactin. The pointed end interaction sites are labelled 1 to 4 [Lau et al., 2021]. (B) An Alphafold prediction of the pointed end complex (Arp11, p25, p27, and p62) and a C-terminal fragment of BICDR (205-394) that includes the Spindly motif. The models are coloured based on the predicted align error (PAE) at Leu347 of BICDR (yellow), with lower values representing higher confidence. The full PAE plot is also shown. (C) Alphafold predictions of cargo adaptors that have been manually linearized such that the coiled-coils are parallel to each other (i.e. each individual model was manually rotated at the disordered loops). White triangles depict breaks in the coiled-coil prediction. The predictions are of full-length proteins unless otherwise stated. LIC-binding motifs (CC1 box, Hook domain, EF hand, RH1 domain), CC2 boxes, and Spindly motifs are coloured according to the legend. Only the CC2 boxes of BICDR1, BICD2, Spindly, TRAK, and Hook3 are shown based on our analyses and previous predictions [Sacristan et al., 2018]. The orientations are C-terminus to N-terminus to match the orientation in other figures. The full PAEs and predicted local distance difference test (pLDDTs) around the Spindly motif can be found in Figure S10. (D) Schematic representation of BICDR-A and BICDR-B showing the relative locations of the interactions with dynein and dynactin (A1 = dynein-A1, A2 = dynein-A2, B1 = dynein-B1, CZ = CapZ, SM = Spindly motif). The black stripes show the site of interaction between BICDR-A and BICDR-B. (E) Schematic showing where the two adaptors bind in Hook3 at the pointed end in comparison to BICDR. Note that Hook3 was predicted to bind site 4 based on mutagenesis studies [Lau et al., 2021].

**Figure S9.**
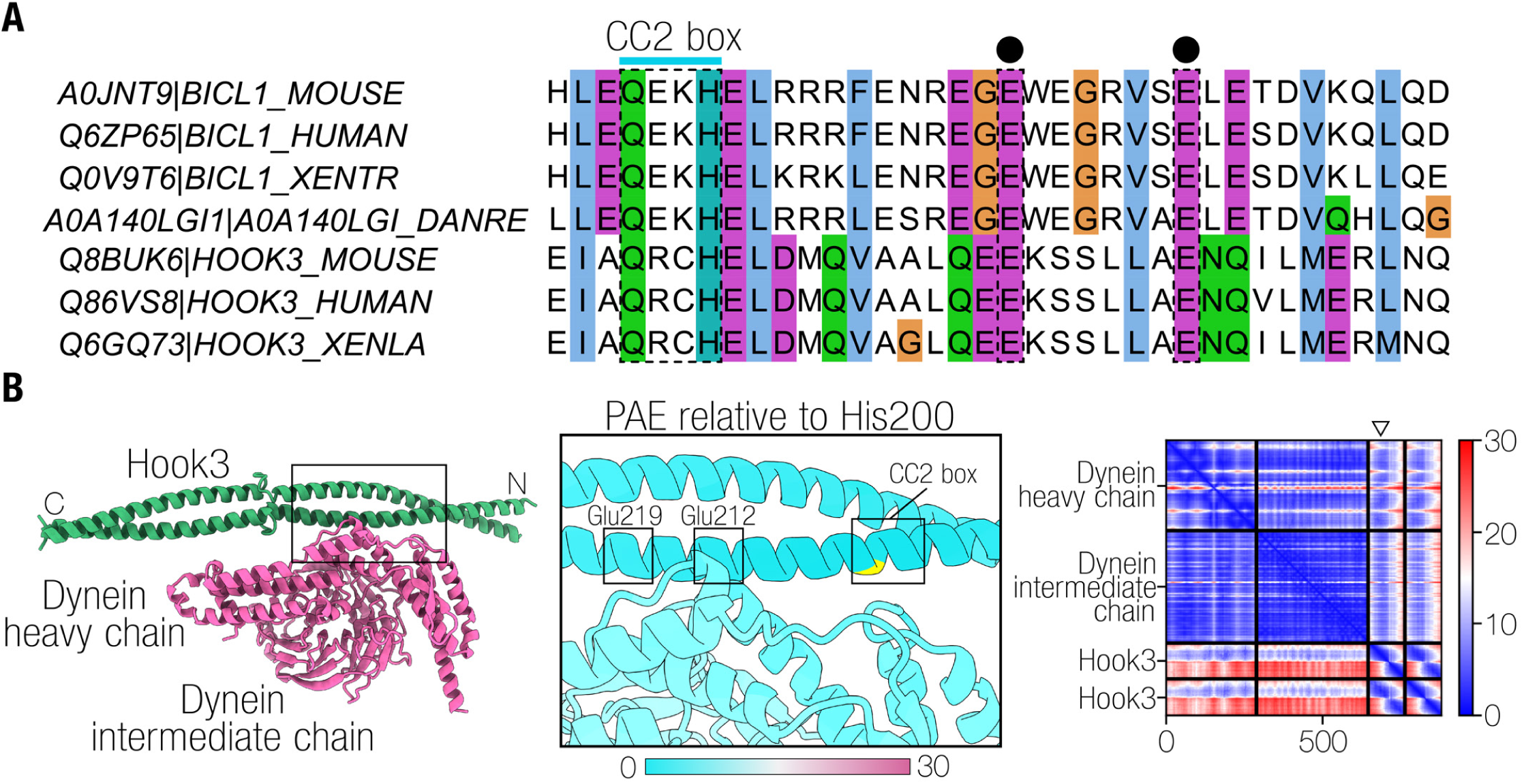
Similarities between the CC2 box of BICDR and Hook3. (A) Alignment of the CC2 boxes of BICDR and Hook family proteins, highlighting the conserved glutamates C-terminal to the motif. The Uniprot codes are indicated on the left. (B) An Alphafold prediction of the Hook3 CC2 box region, containing two copies of Hook3 (fragment 172-287), dynein heavy chain (fragment 576-864), and dynein intermediate chain (fragment 226-583). In the middle, the PAE is displayed on the models relative to His200 (yellow), with lower values representing higher confidence. The full PAE plot is shown on the right.

**Figure S10.**
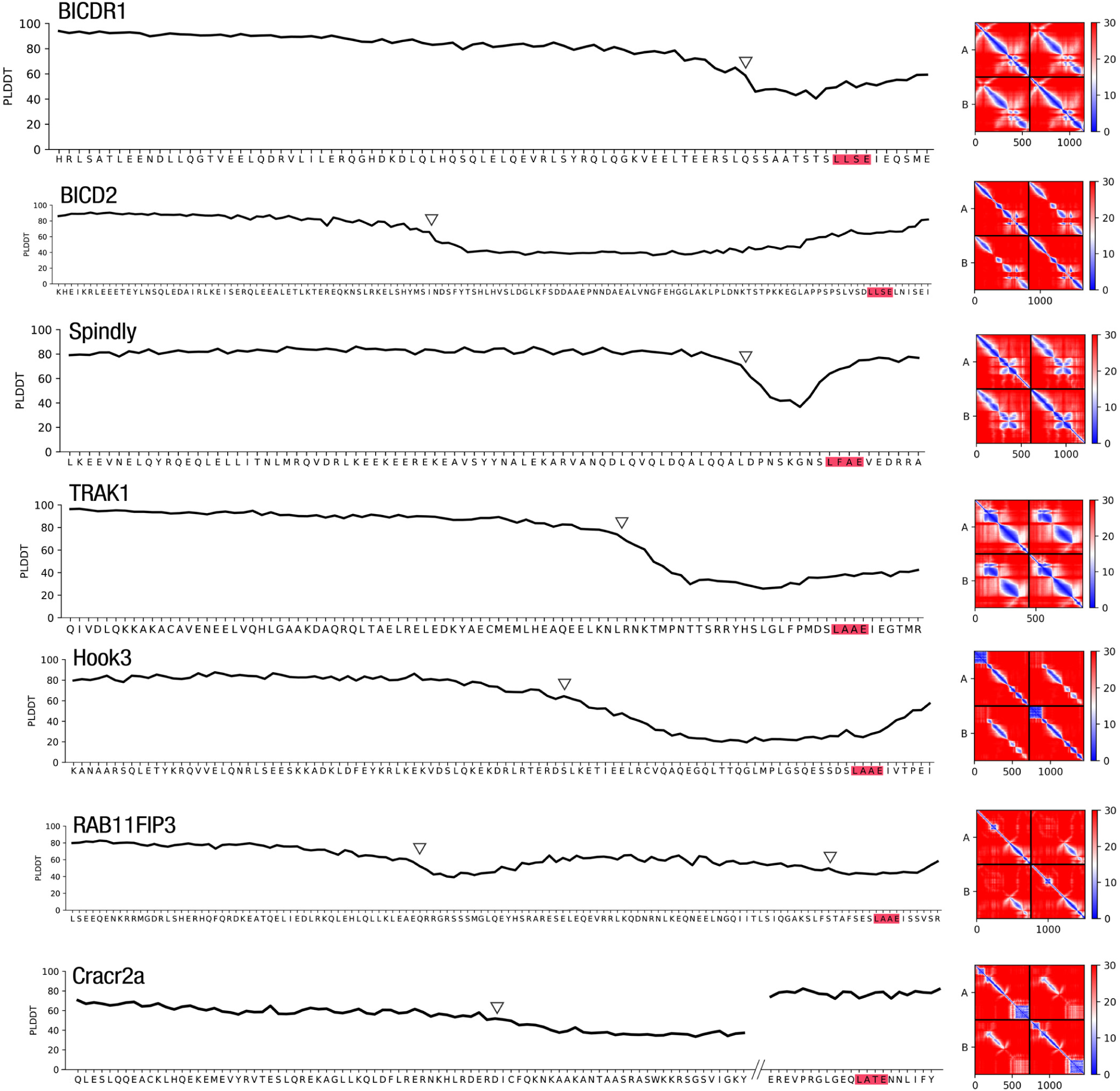
Cargo adaptor prediction confidence plots. The pLDDTs from one chain of each of the cargo adaptors around the Spindly motif are shown in the plots on the left. There is a consistent drop in accuracy preceding the motif, indicating a higher likelihood of disorder. The white triangles point to breaks in the coiled-coil secondary structures in the resulting predictions. The PAE plots are shown on the right.

